# Stochastic model of T Cell repolarization during target elimination (I)

**DOI:** 10.1101/822171

**Authors:** I. Hornak, H. Rieger

## Abstract

Cytotoxic T lymphocytes (T) and natural killer (NK) cells are the main cytotoxic killer cells of the human body to eliminate pathogen-infected or tumorigenic cells (i.e. target cells). Once a NK or T cell has identified a target cell, they form a tight contact zone, the immunological synapse (IS). One then observes a re-polarization of the cell involving the rotation of the microtubule (MT) half-spindle and a movement of the microtubule organizing center (MTOC) to a position that is just underneath the plasma membrane at the center of the IS. Concomitantly a massive relocation of organelles attached to MTs is observed, including the Golgi apparatus, lytic granules and mitochondria. Since the mechanism of this relocation is still elusive we devise a theoretical model for the molecular motor driven motion of the MT half-spindle confined between plasma membrane and nucleus during T cell polarization. We analyze different scenarios currently discussed in the literature, the cortical sliding and the capture-shrinkage mechanisms, and compare quantitative predictions about the spatio-temporal evolution of MTOC position and spindle morphology with experimental observations. The model predicts the experimentally observed biphasic nature of the repositioning process due to an interplay between spindle geometry and motor forces and confirms the dominance of the capture-shrinkage over the cortical sliding mechanism when MTOC and IS are initially diametrically opposed. We also find that the two mechanisms act synergetically, thereby reducing the resources necessary for repositioning. Moreover, it turns out that the localization of dyneins in the pSMAC facilitates their interaction with the MTs. Our model also opens a way to infer details of the dynein distribution from the experimentally observed features of the MT half-spindle dynamics. In a subsequent publication we will address the issue of general initial configurations and situations in which the T cell established two immunological synapses.

## 1 Introduction

Cytotoxic T lymphocytes and natural killer cells have a key role in our immune system by finding and destruction of virus-infected and tumor cells, parasites and foreign invaders. Once a T-Cell leaves the thymus, it circulates through the organism in search of antigen-presenting cell (APC). The directional killing of an APC is completed in three subsequent steps. First, T cell receptors (TCR) bind to antigens on the surface of the APC presented by the major histocompatibility complex (1–6) leading to the creation of a tight contact zone called immunological synapse (IS) (7–9) composed of multiple supramolecular activation clusters (9–11). In the second step, the cell repolarizes by relocating the microtubule organizing center (MTOC) towards the IS (12–18) under the influence of forces exerted on MT. Moreover, since mitochondria, Golgi apparatus and Endoplasmic reticulum are attached to MTs, these organelles are dragged along with the spindle (16, 19–24). Consequently, the repolarization process involves massive rearrangements of the internal, MT associated structure of the cell. In the third step, the T-Cell releases at the IS cytotoxic material (e.g. the pore forming protein perforin and the apoptosis inducing granzyme) from vesicles, the lytic granules, (25–29) towards the APC leading to its destruction (29–37). Hence, the repositioning of the MTOC plays a critical role in the elimination of the target cell since it transports the lytic granules to the IS (38, 39).

The IS is partitioned into several supramolecular activation clusters (SMACs) including the peripheral SMAC (pSMAC) and the central SMAC (cSMAC) (7, 9, 11, 40, 41), in which TCR (cSMAC), adhesion molecules AND actin are organized. Dynein, a minus-end directed (towards the MTOC) molecular motor protein, is absolutely necessary for the repolarization to take place as was experimentally demonstrated with knock-out experiments (42). Once the T-cell is activated the adaptor protein ADAP forms a ring at the periphery of IS with which dynein co-localizes (43, 44). Concerning the underlying mechanism, it was proposed that the repolarization is driven by the cortical sliding mechanism (43, 45) in which dyneins anchored on the cell membrane step on the MT towards the minus-end and thus pull the centrosome towards the IS. The first experimental indications for cortical sliding came from the observation of the cytoskeleton movement using polarization light microscopy (17). Subsequent experiments indicate that the IS periphery, in particular the ring shaped pSMAC, is the region where dyneins attach to and pull on MTs (17, 43).

Repositioning was observed in various experiments. Focused activation of photo-activable peptide-MHC on the glass surface was used in (46). In (16) the repositioning was observed alongside with the rotation of the mitochondria, which provided evidence that the mitochondria are dragged with the spindle. Detailed observations were made by Yi et al (14) providing a new insight into the mechanism of repolarization. In (14) an optical trap was used to place an APC so that the initial point of contact is in diametrical opposition to the current position of the MTOC, which allowed for dynamical imagining in a quantitative fashion. During the experiment, the deformations and changes in MT structures were observed and the position of the MTOC tracked. First of all, Yi et al. (14) provided strong experimental evidence against a cortical sliding mechanism. Instead the observations indicate that the MTOC is driven by a capture-shrinkage mechanism (47) localized in a narrow central region of the IS. The capture-shrinkage mechanism involves dynein that interacts in an end-on fashion with the plus-end of a MT which is fixed in a position on the membrane of the cell where the MT depolymerizes. The MT shrinkage part happens plausibly because dynein pulls the MT plus-end against the cell membrane, which increases the force dependent MT depolymerization rate (47).

In sequences of microscope pictures (14) showed that MTs reach from the MTOC to the IS and bend alongside the cell membrane. Subsequently, MTs form a narrow stalk connecting the MTOC with the center of IS. The plus-end of the MTs in the stalk, while captured in the center of the IS, straighten (probably under tension due to the dynein pulling at the plus-end) and shrink by depolymerization at the capture point. Consequently, the MTOC is dragged towards the center of IS, which invaginates into the cell, further proving the location of the main pulling force. When the MT depolymerization was inhibited by taxol, the MTOC repositioning slowed down substantially. These observations supported the hypothesis that the capture-shrinkage mechanism plays a major role. Additionally, Yi et all (14) reported that the repositioning is biphasic and that the two phases differ in the speed of the MTOC and the orientation of its movement. In the first, so-called polarization phase, the MTOC travels quickly around the nucleus of the cell in a circular manner. The polarization phase ends when the MTOC is approximately 2*μ*m from the center of IS. Subsequently, during the second “docking phase” the MTOC travels directly towards the IS with substantially decreased speed.

The cortical sliding mechanism alone was previously analyzed with a deterministic mechanical model (48), where it was demonstrated that mechanism is capable of reorienting the MTOC into a position under the IS underneath certain conditions. Furthermore, oscillations between two IS were studied in different situations. Nevertheless, the forces in the model were deterministic, neglecting the stochastic nature of dynein attachment, detachment and stepping, leaving various experimental observations unexplained, as for instance the preferential attachment of MTs to dynein anchored in the periphery of the IS.

Sakar etal (49) hypothesized that dynamic MTs find the central region of the IS, where they can be captured by dynein by growing from the MTOC in random directions, analogous to the search and capture mechanism during the formation of the mitotic spindle. Once MTs attach to dynein in the central region of the IS relocation of the MTOC starts, which is the process that is analyzed in this paper.

In spite of these detailed experimental observations many aspects of the internal mechanisms driving the relocation of the MTOC during the T-Cell repolarization remain poorly understood like the cause of the transition from the polarization to the docking phase. Yi et al argue that a resistive force emerges when the MTOC-IS distance is around 2*μ*m leading to a reduction in the MTOC velocity. The potential causes are physical impediments to MTOC translation or a reduced attachment or force development of molecular motors. Moreover, the experiments of Yi etal were performed with specific initial positions of the IS and the MTOC, being diametrically opposed. The question arises whether the observed dominance of the capture-shrinkage mechanism would be robust in naturally occurring situations in which the initial position of the MTOC is not in diametrical opposition to the IS. If capture-shrinkage is the truly dominant mechanism, what is the role of cortical sliding? Finally, why are cortically sliding MTs caught just on the periphery of the IS (17), is it caused purely by the colocalization of dynein with ADAP ring (43)? The answers to these question is still elusive and in this work we analyze them in the framework of a quantitative theoretical model for the relocation of the MTOC after IS formation.

We distribute our analysis into two consecutive publications. In this, first, publication we describe the theoretical model we use and present our results focusing on the experiments described in (14), (17) and (43) and on an analysis of the two mechanisms: cortical sliding and capture shrinkage. This comprises the setup in which the T-cell has one IS and the intitial positions of IS and MTOC are diametrically opposed to each other.

A subsequent, second, publication will focus on quantitative predictions of our model for situations that have not yet been analyzed experimentally. There we will focus on the repolarization following initial configurations not realized in (43), which will also provide additional insight into the different effects of the two mechanisms, cortical sliding and capture shrinkage. Moreover we will analyze the, eventually oscillating, MT/MTOC movement without IS and with two IS.

## 2 Methods

### 2.1 Computational model

The cell and its nucleus are modeled as two concentric spheres of radius 5*μ*m and 3.8*μ*m, respectively. The model of the cytoskeleton consists of microtubules (MT) and the microtubule organizing center (MTOC) 1. MTs are thin filaments with a diameter of ca 25nm (50–52). The measured values of flexural rigidity varies between experiments (53, 54), in our model we take 2.2 * 10^−23^ Nm^2^ (55) yielding a persistence length larger than 5mm exceeding the size of the cell by three orders of magnitude.

A single MT is represented by a bead-rod model (56). Since repolarization occurs on a time scale of seconds, the growth of MTs is neglected. The beads move under the influence of forces to be described below (and defined in detail in the SI 1a): bending, drag, molecular motor and stochastic forces. Assuming zero longitudinal elasticity of the MTs we use constrained Langevin dynamics to model the motion of the MTs, see SI. Repulsive forces acting on the MT segments confine the cytoskeleton between the nucleus and the cell membrane.

The MT organizing center (MTOC) is a large protein complex which has a complex structure composing of mother and daughter centrioles (57–60) embedded in the pericentriolar material(PCM) (61–63). MTs nucleate from gamma-tubulin-containing ring structures within PCM mainly at the appendages of mother centriole (57, 64). MT can sprout from the MTOC in all directions. MTs whose original direction is approximately parallel to the membrane of the cell, will continue to grow to the cell periphery. Other MTs will soon hit the wall of the cell or its nucleus. Such MT can either bend and assume new direction parallel to the cell membrane, or undergo MT catastrophe. Therefore, the long MTs are seemingly always sprouting from the MTOC in one plane, as can be seen in (14). Consequently, we model the MTOC as a planar, rigid, polygon structure Figure (SI 1b)) from which MTs emanate in random directions by fixing the positions and directions of their first segment, (SI 1b)). MTs sprout from the MTOC to the cell periphery 1a.

Unattached dynein is represented just with one point on the surface of the cell. If the dynein is closer to the MT than *L*_0_, protein attaches with a probability *p_a_*. Dynein motors are distributed randomly in specific,spatially varying concentrations, on the cell boundary. Attached dynein is represented by an fixed anchor point located on the cell boundary and an attachment point located on a MT, both being connected by an elastic stalk of a length *L*_0_ (65, 66). The force exerted on a MT 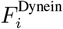 depends on stalks elastic modulus *k*_Dynein_ and prolongation. Dynein stepping depends on the magnitude of the force and its orientation. If the force is parallel to the preferred direction of the stepping, attachment point moves one step to the MT minus-end (towards the MTOC) with constant probability *p*_−_. If the orientation of the force is opposite and its magnitude smaller than stall force *F_S_*, dynein makes one step towards to the minus-end with force-depending probability. If 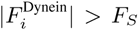 and the force has unfavorable direction, dynein makes one step to the plus-end with constant probability *p*_+_. The steps of dynein have varying lengths(66) but for simplicity we set it to the most frequently measured value of *d*_step_ = 8*n*m. The probability of detachment, *p*_detach_, increases with force.

Experimentally two mechanisms by which dynein act on MTs have been identified: cortical sliding (17), where MTs under the effect of dynein move tangentially along the membrane, and capture shrinkage (14), by which MTs under the effect of dynein are reeled in towards the membrane and concomitantly depolymerized (sketched in Figs. 1)d and 1)e.

**Figure 1:**
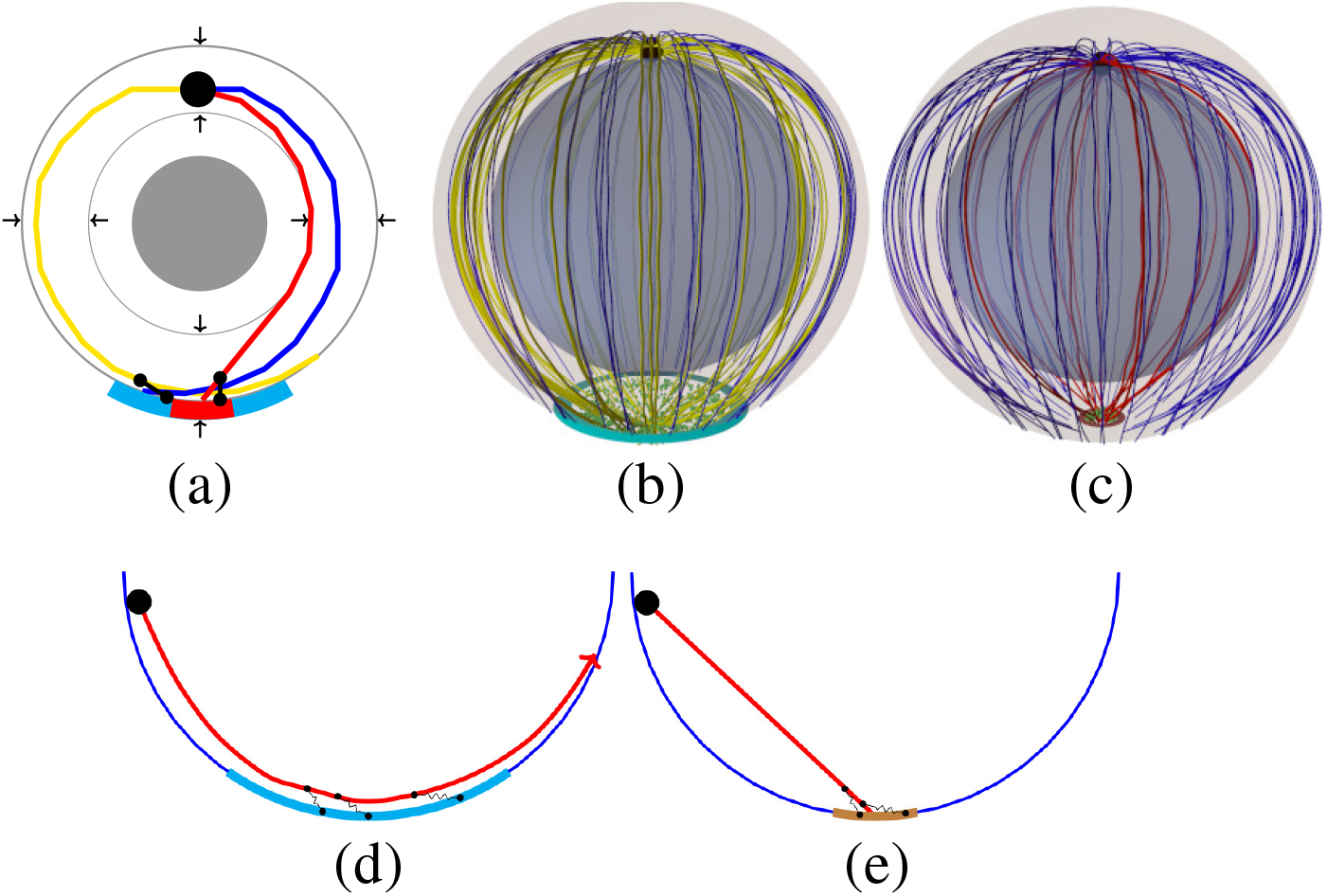
(a)(b)(c) Sketch of the model: The large black sphere represents the MTOC, MTs are represented as lines in three different colors, MTs not attached to dynein (blue), attached to cortical sliding dynein (yellow) and attached to capture-shrinkage dyneins (red). (a) Two-dimensoinal cross-section of the model: The gray circle indicates the center of the nucleus, black arrows the repulsive forces acting on MTs segments, small black dots represent the anchor point of dyneins. (b)(c) Three-dimensiomnal sketch of of the cell model. The outer transparent and inner spheres represent the cell membrane and the nucleus of the cell, respectively. (b) blue disk represents the IS, where cortical sliding dynein is anchored. Small green dots in the IS represent randomly distributed dynein. (c) Brown disk represents the central region of the IS where capture-shrinkage dynein is anchored. (d)(e) Sketch of the cortical sliding mechanism (d) and the capture-skrinkage mechanism (e). Large black circle: MTOC, small black dots on the membrane: dynein anchor points, small black dots on the MTs: attachment points, cyan membrane segment: IS, brown membrane segment: central region of the IS where capture-skrinkage dynein is anchored. Note that MT depolymerize when pulled by capture-shrinkage dynein towards the membrane.

The IS is divided into two regions: the center, where dyneins act on MTs via the capture shrinkage mechanism (14) and the complete IS, where dyneins act via the cortical sliding mechanism. Each region is modeled as an intersection of the cell sphere with a cylinder, 1, with radius *R*_IS_ = 2*μ*m for the complete IS and *R*_CIS_ = 0.4*μ*m for the central region. Dyneins are distributed randomly with uniform area density *ρ*_IS_ in the small central region, denoted as capture-shrinkage dynein, and density 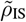 in the larger region of the whole IS, denoted as cortical-sliding dynein.

## 3 Results

We analyzed the role of the cortical sliding and capture shrinkage mechanisms during repolarization by computer simulations of the model defined in the previous section. Each simulation run is initialized with the mechanical equilibrium (minimum elastic energy) configuration of the MT/MTOC-system and all dyneins bein detached. During the integration of the equation of moition various quantities are calculated: distance between centers of the MTOC and the IS, *d*_MIS_, number of dyneins attached to the MTs, *N*_dm_, velocity of the MTOC, *υ*_MTOC_, distance between the MTOC and the center of the cell, *d*_MC_. For each point in the parameter space these quantities were averaged over 500 simuation runs. Results are shown with the standard deviation as error bars only when they are larger than the symbol size.

### 3.1 Capture-Shrinkage mechanism

The repositioning process under the effect of the capture-shrinkage mechanism is visualized in Fig. 2. In Figs. 2a and 2d, it can be seen that initially the attached MTs aim from the MTOC in all directions. Subsequently, the stalk of MTs is almost formed in the middle phase of the repositioning Fig. 2b and Fig. 2e, and it is fully formed as the MTOC approaches the IS, see Fig. 2c and 2f and the supplementary movie Movie_1 showing the time-evolution of the spindle configuration under the effect of the capture-shrinkage mechanism with 100 MTs and dynein density *ρ*_IS_ = 100*μ*m^−2^.

**Figure 2:**
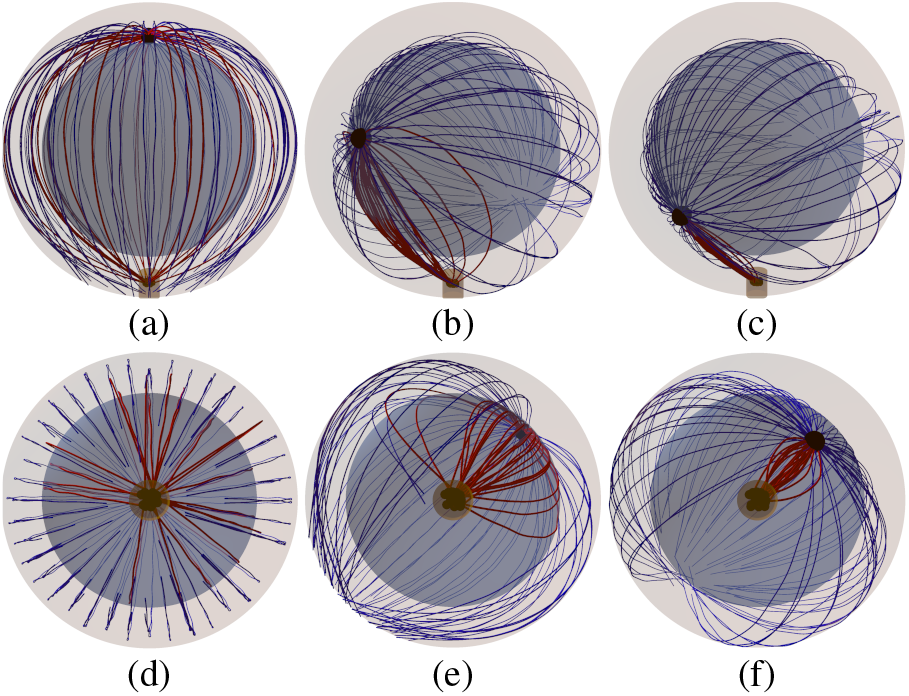
Snapshots from the time-evolution of the spindle configuration under the effect of the captureshrinkage mechanism alone (dynein density *ρ*_IS_ = 100*μ*m^−2^) from horizontal (upper row) and vertical perspective. MTs are connected to the MTOC indicated by the large black sphere. Blue and red curves are unattached and attached MTs. Small black spheres in the IS represent dyneins. The brown cylinder indicates the center of the IS, where the capture-shrinkage dyneins are located. (a)(d) *t* = 20s, *d*_MIS_ = 9*μ*m. Initially the attached MTs sprout from the MTOC in all directions directions. (b)(e) *t* = 65s, *d*_MIS_ = 6*μ*m. As time progresses, microtubules form a stalk connecting the MTOC and the IS. (c)(f) *t* = 80s, *d*_MIS_ = 2.5*μ*m. The stalk is fully formed and it shortens as the MTOC approaches the IS.

The process can be divided into three phases based on the time-evolution of the MTOC velocity, see Fig. 3b. In the first phase, when the distance between MTOC and the center of the IS is 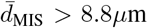, the velocity changes rapidly in the first seconds of the process and then falls to a local minimum. In the second phase, the velocity continually increases to a maximum and then in the third phase, it decreases again. By comparison of Fig. 3b and Fig. 3c, it can can be seen that the time-evolution of the velocity corresponds to the time-evolution of the number of dyneins acting on MTs. The evolution of the number of attached dyneins during the first phase can be understood from an analysis of Figs. 3d, 2a and 2b. At the beginning of the simulations, a substantial number of MTs intersects the IS (visually demonstrated in Fig. 2a and Fig. 2d) resulting in a fast increase of the number of attached dyneins. Since the MTs attached to dynein sprout from the MTOC in every direction, c.f. Fig. 3e, the MTOC moves towards the IS and, simultaneously, to the nucleus of the cell, see Fig. 3d. As the MTOC approaches the nucleus of the cell, the nucleus starts to oppose the movement by repelling the MTs and, at the end of the first phase, the MTOC. Therefore, as the pulling force of the dyneins is opposed by the nucleus, the dyneins detach since the detachment rate is force-dependent.

**Figure 3:**
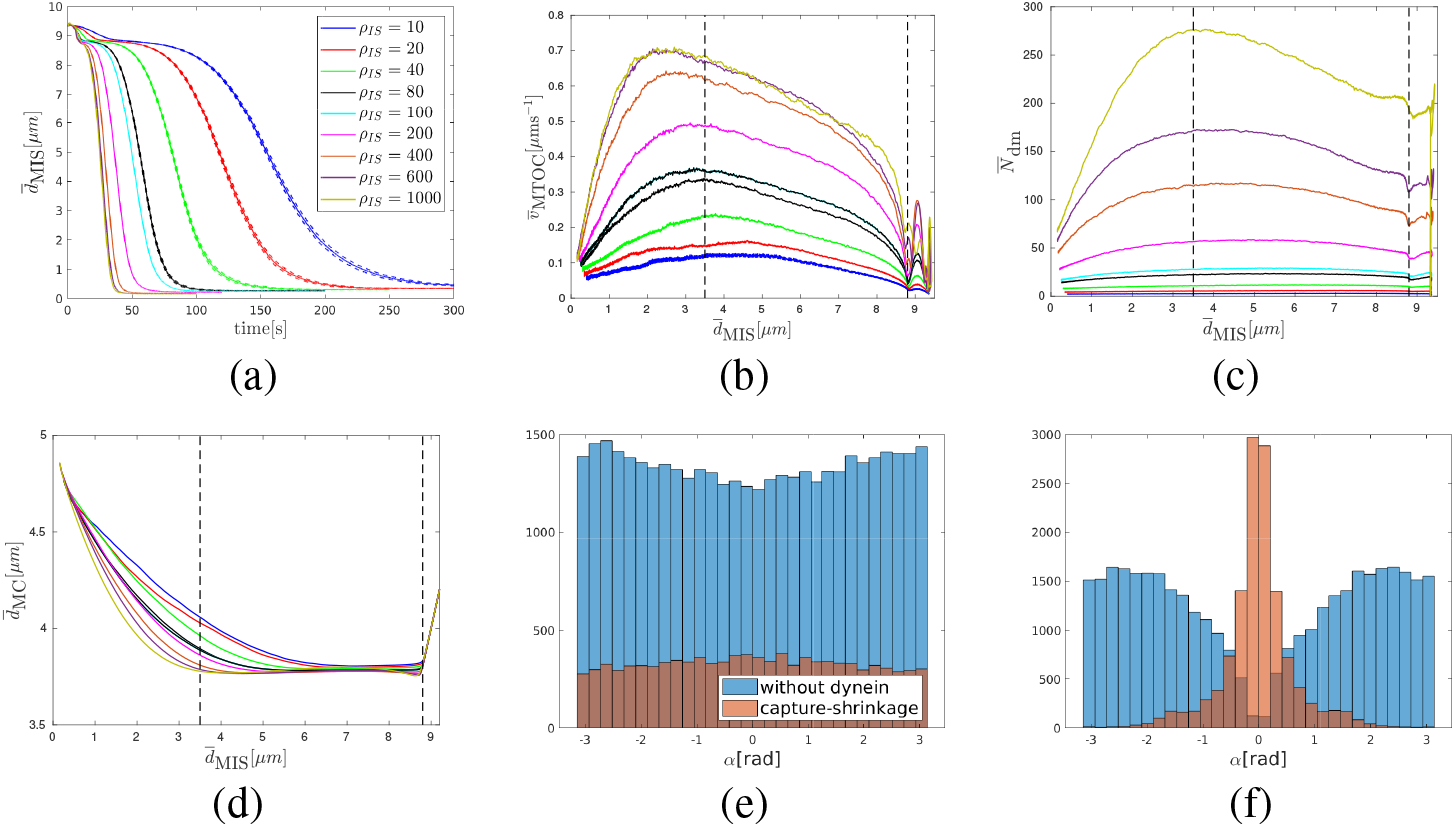
Capture-shrinkage mechanism: (a) Dependence of the average MTOC-IS distance 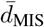 on time. (b)(c)(d) Dependencies of the average MTOC velocity 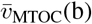, the number of dyneins acting on microtubules 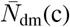 and the MTOC-center distance 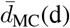 on the average MTOC-IS distance. Black dashed lines denote transitions between different phases of the re-positioning process. (e)(f) Histograms of the angles between the first MT segments and the direction of the MTOC movement for a dynein density *ρ*_IS_ = 100*μ*m^−2^ (e) *t* = 1s, 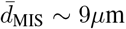. (f) *t* = 60s, 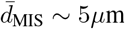.

The increase of the number of attached dyneins 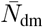 in the second phase can be explained by considering the fact that the MTOC slides over the surface of the nucleus and the MT stalk forms. At the beginning, the nucleus presents an obstacle between the MTOC and the IS, see Fig. 2a. The opposing force from the nucleus decreases with the approach of the MTOC towards IS. At the end of the repositioning, the nucleus does not longer stands between the two objects, see Fig. 2c and 2f. Therefore, the opposing force from the nucleus contributing to dynein detachment decreases. More importantly, attached MTs are form the MT stalk. The angle *α* between the first segment of the MT and the direction of the MTOC movement is used to describe the deformation of the cytoskeleton structure and stalk formation. At the beginning of simulation(the first phase and the beginning of the second), attached MTs aim in every direction, see Fig. 3e, visualized in Fig. 2a and 2d. Therefore, the dyneins pull in multiple directions which makes them oppose each other leading to dynein detachment. After a few seconds, the MTOC travels in the direction of the biggest pulling force. Consequently, the attached MTs form a stalk as the simulation progress and dyneins act in alignment, see Figs. 3f, 2b and 2e. They do not longer oppose each other, but they share the load from opposing forces. Consequently, the detachment probability of dynein decreases with the opposing force and the number of attached dyneins increases.

The number of dyneins decreases in the final phase when 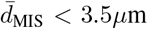, see Fig. 3c. Unattached MTs in the IS are pushed backward by viscous drag as the MTOC moves to the IS. As a result, one observes an “opening” of the MT spindle, c.f. Figs. 3e, 2c and 2f. Unattached MTs do not intersect the IS, see Fig. 3f, and cannot attach to dynein. The attached MTs shorten due to depolymerization further lowering the probability of dynein attachment. Moreover, an opposing force arises from the cytoskeleton being dragged from the nucleus to the membrane, see Fig. 3d.

To summarize, the trajectory of the MTOC towards the IS displays three phases, where the two longer phases hae been reported also in the experiment (14) but not the short initial phase. First the MTOC descends to the nucleus, see Figs. 3a and 3d, then it moves to the IS fast and then slows down during the last 2*μ*m, see Fig. 3b. Once the MTOC bypasses the nucleus, it moves away switching from purely circular to partially radial movement, see Fig. 3d. The variation of the MTOC velocity, its modulus and its direction, is clearly visible in the supplementary movie Movie_2, showing a simulation with a smaller nucleus radius *r_N_* = 3.3*μ*m. Note that the duration of the complete repositioning process in the experiments is ca 60-90 sec, which our model predicts to be achieved by a dynein density of 80-200 *μ*m^−2^.

### 3.2 Cortical Sliding mechanism

For low, medium and high densities one observes for each a different characteristic behavior. In the regime of low dynein densities, 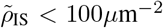, the re-polarization speed increases with the dynein density and the MTOC moves directly to the IS, see Fig. 5a. For medium dynein densities, 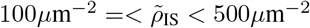, the MTOC movement is more complex, see Fig. 7a. For high dynein densities 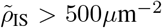 the re-polarization speedsurprisingly decreases with 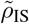, see Fig. 8a.

#### 3.2.1 Cortical sliding with low dynein densities

The supplementary movie Movie_3 shows the MTOC repositioning under the effect of the cortical-sliding mechanism with 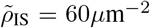. It shows MTs sprouting in all directions in the initial stage, the subsequent stalk formation and the final slowing down of the MTOC.

In Fig. 5b the dependence of the MTOC velocity on MTOC-IS distance is shown. As in the case of the capture-shrinkage mechanism, the time-evolution of the MTOC velocity can be divided into three phases. However, the transition points between the second and the third phase depend on the density 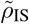. Similarly to the case of the capture-shrinkage mechanism, the behavior in the first phase can be explained by the interplay of fast attaching dyneins and forces from the nucleus. In the second phase, the velocity of the MTOC increases in spite of a continuously decreasing number of attached dyneins, see Figs. 5b and 5c, which is due to the alignment of the MTs. The explanation the alignment of MTs. Initially, attached MTs aim in all directions, see Fig. 5e, as for the capture-shrinkage mechanism, c.f. 4a and 4d. Consequently, MTs, whose original orientation does not correspond to the movement of the MTOC, detach from dynein, see Figs. 5f, 4b and 4e. The histogram in the intermediate state of repositioning 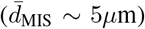 shows that attached MTs are aligned and less in numbers. The MTOC does not significantly recede from the nucleus at the end of repositioning, see Fig. 5d, which implies that the MTs do not follow the cell membrane (with the capture-shrinkage mechanism MTs always touch the membrane), see Figs. 4c and 4f. Consequently, the attachment probability is lower and leads to the decrease in speed in the third phase.

**Figure 4:**
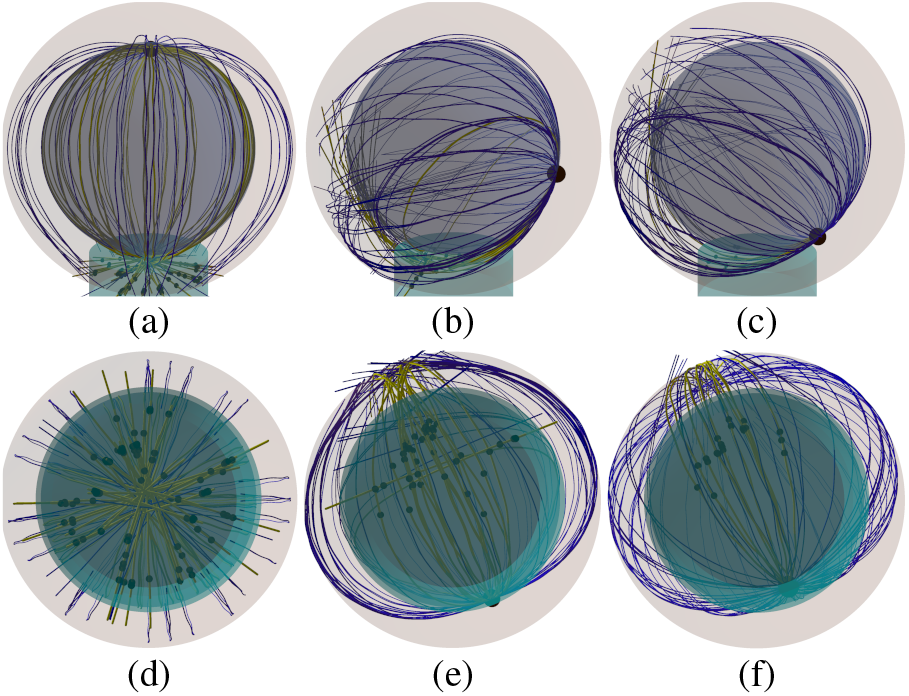
Snapshots from the time-evolution of the spindle configuration under the effect of cortical sliding mechanism with a low dynein density, 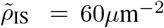, from two perspectives. The cyan cylinder indicates the IS area. Blue and yellow lines are unattached and attached MTs, respectively. The black spheres in the IS are the positions of dynein attached to MTs. (a)(d)*t* = 5s, 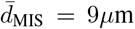. Originally, the attached microtubules aim from the MTOC to every direction. (b)(e)*t* = 55s, 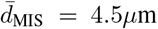. MTs attached to dynein aim predominantly in one direction. (c)(f)*t* = 90s, 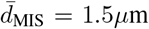. Just a few MTs remain under the actions of cortical sliding and they rarely touch the surface of the cell in IS.

**Figure 5:**
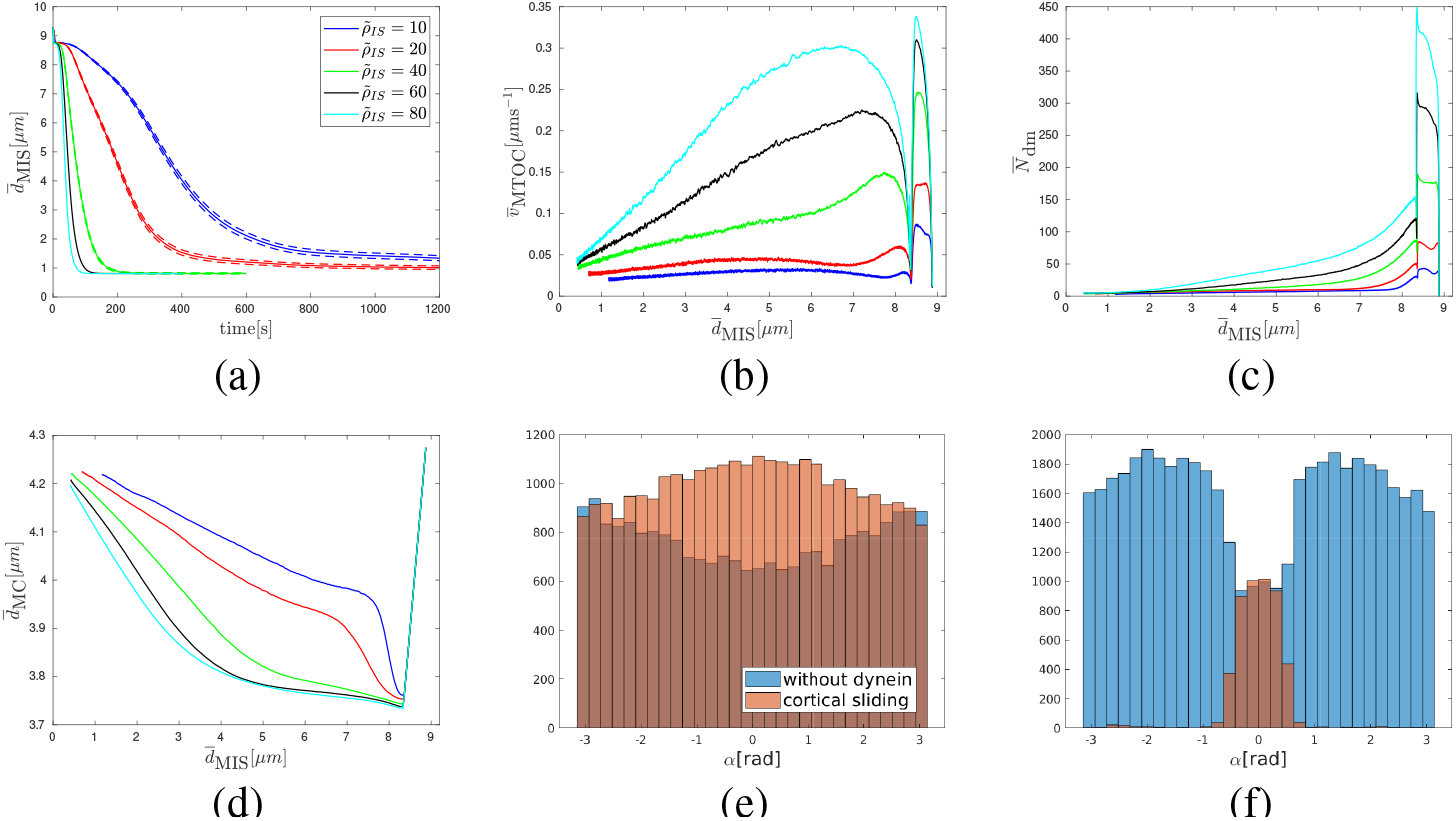
Cortical sliding with low dynein densities 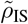. (a) Dependence of the average MTOC-IS distance 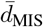 on time. (b)(c)(d) Dependencies of the average MTOC velocity 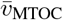 (b), number of dyneins acting on MTs 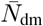 (c) and MTOC-center distance 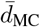 (d) on the average MTOC-IS distance. (e)(f) Histogram of the angles *α* between the first MT segment and the MTOC motion. From 500 simulation runs for dynein density 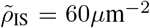. (e) *t* = 5s, 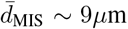. (f) *t* = 65s, 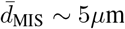.

#### 3.2.2 Cortical sliding with medium dynein densities

The differences between the behavior with low and medium dynein densities for the cortical sliding mechanism are analyzed in this section. The supplementary movie Movie_4 shows the repositioning with 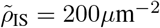. The repositioning is very fast and the MT spindle is considerably deformed. Moreover, the MTOC passes the IS and subsequently returns to the center of IS. Five seconds after the initialization MTs in all directions are attached, see Fig. 6a and 6d, but the direction of the MTOC motion is already established, see Fig. 7b. Contrary to the case of low densities, the dynein forces are sufficiently strong to hold attached MTs. Subsequently some MTs do not detach but take a direction partially aligned to the MTOC movement, see Fig. 7c and 7d. These filaments follow a trajectory between the MTOC and the dyneins that is given by its length and tendency to minimize bending energy. MTs aligned with the MTOC movement slide the fastest. Logically, the majority of MTs have a length longer than the MTOC-IS distance. Therefore, they cannot fully align with the MTOC movement. Subsequently, the maximum of the histogram is not at |*α*| ~ 0.0*π* but at |*α*| ~ 0.6*π*, see Fig. 7c and 7d and the cytoskeleton is deformed, c.f. Figs. 6c and 6e. Here it can also be noticed that the MTs with sprouting orientation opposite to the MTOC movement are held by dynein, which provides further testimony of its strength. Nevertheless, the large majority of MTs are aligned with the MTOC movement, which is the reason for its speed. It can be seen from comparison between Figs. 5a, 7a and 8a and from Fig. 9a and 9b that the velocity of the MTOC with medium densities of cortical sliding dynein is unparalleled. The strength of dynein and twisting of MTs causes large deformations of the MT spindle, compare Figs. 4b and 4e with Figs. 6b and 6e.

**Figure 6:**
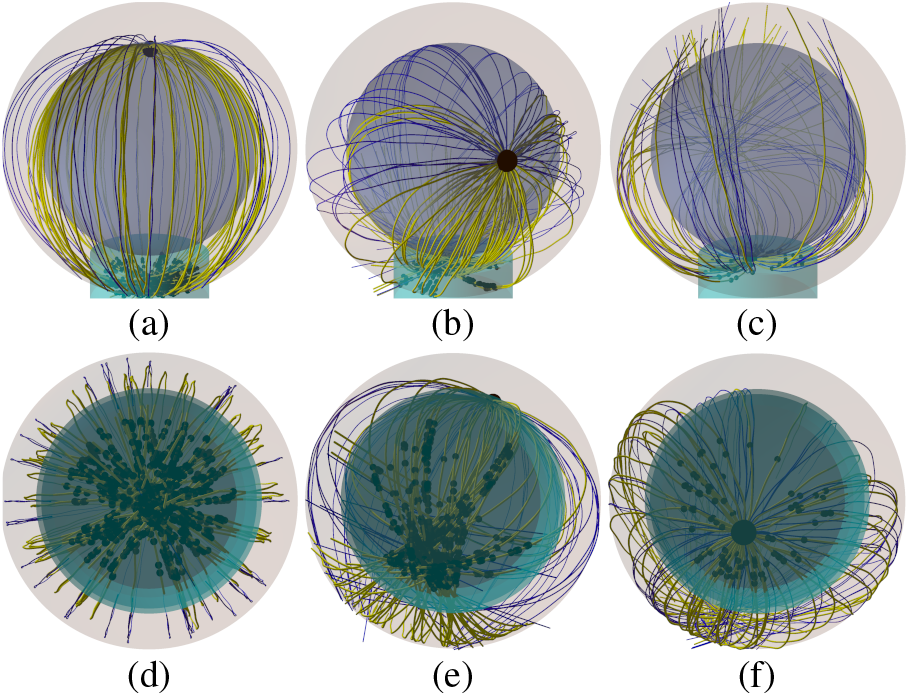
Snapshots from the time-evolution of the spindle configuration under the effect of cortical sliding alone, with a medium area density of dynein 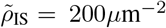) from two perspectives. (a)(c) *t* = 5s, 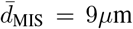. MTs sprout from the MTOC in all directions. (b) (d) *t* = 15s, 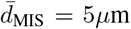. The majority of MTs is attached and the MT spindle is deformed. (c) (e) *t* = 25s, 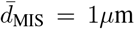. At the end of the repositioning, the MTOC passed the center of the IS and attached MTs aim in all directions.

**Figure 7:**
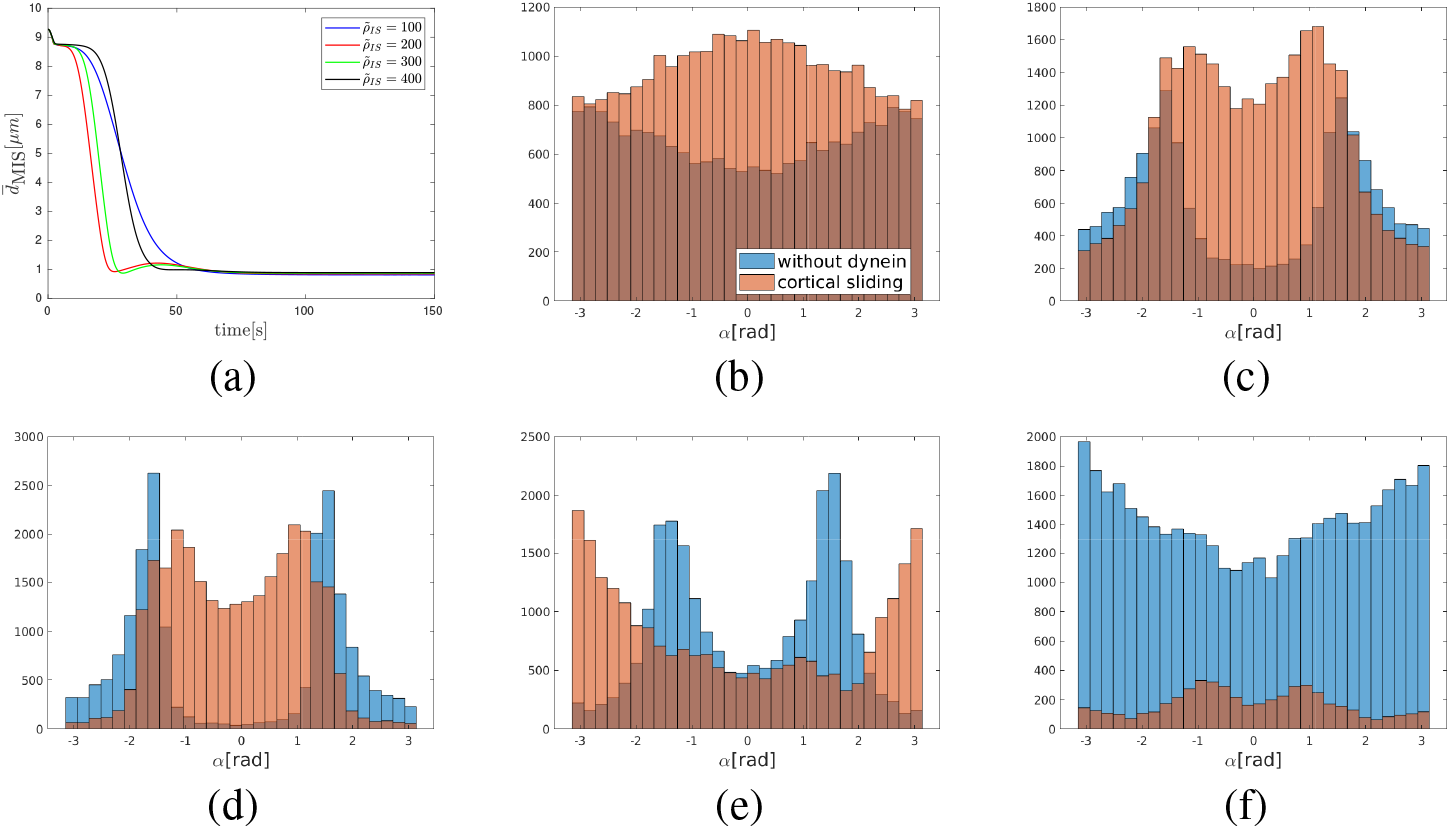
Cortical sliding mechanism with medium dynein densities 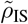. (a) The dependence of the MTOC-IS distance 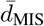 on time. Histogram of the angles *α* between the first MT segment and the MTOC motion from 500 runs of simulation for the case of area density of cortical sliding dyneins 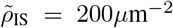. (b)*t* = 5s, 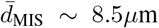. (c)*t* = 15s, 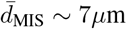. (d)*t* = 20s, 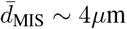. (e)*t* = 25s, 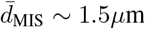, other side of IS. (f)*t* = 60s, 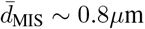.

**Figure 8:**
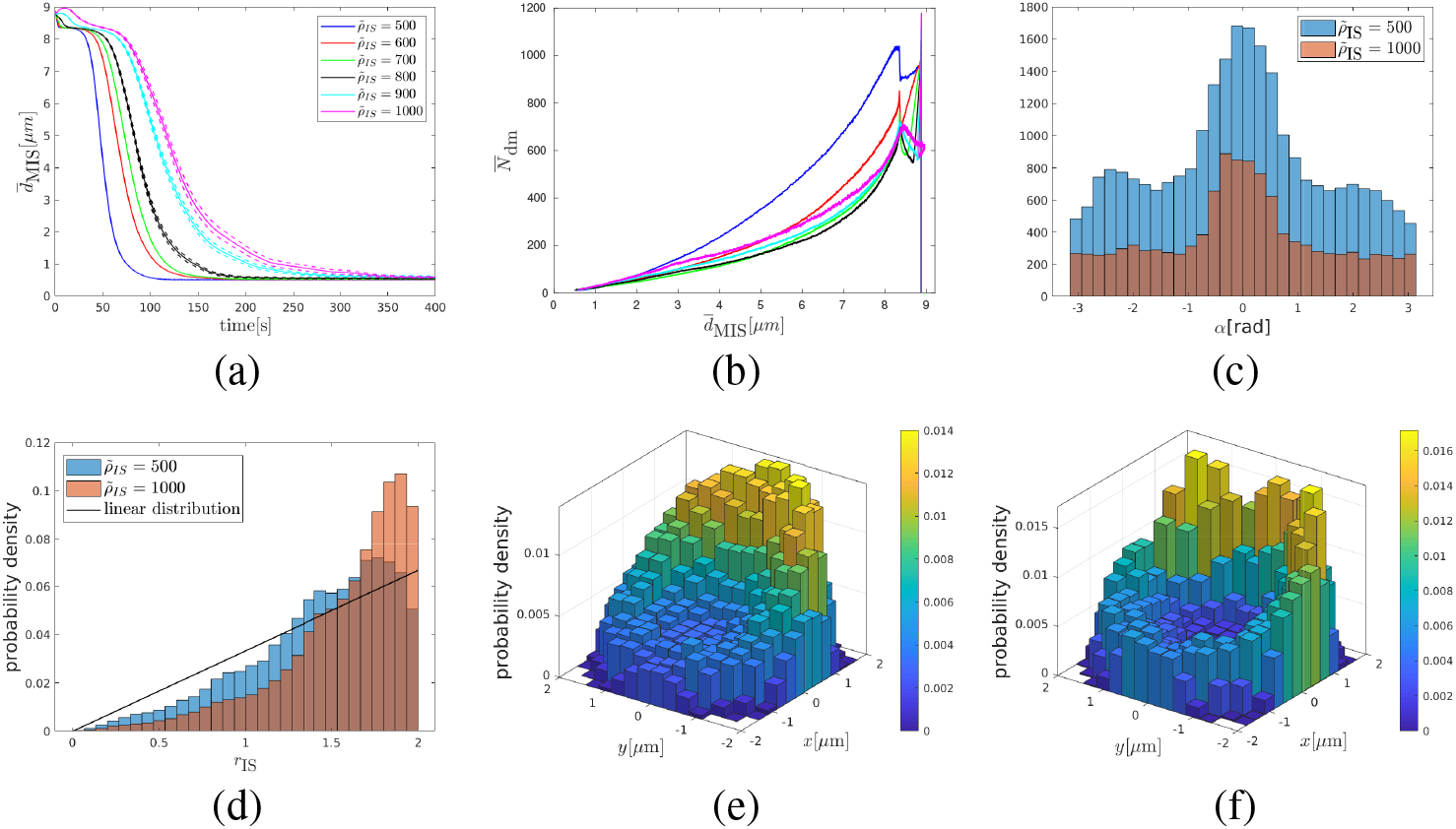
Cortical sliding with high dynein densities 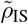. (a) The dependence of the average MTOC-IS distance 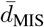 on time. (b) The dependence of average number of dyneins 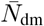 on the MTOC-IS distance. (c) Histogram of the angle *α* between the first MT segment and the direction of the MTOC motion. From 500 simulation runs, 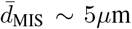. (d) Probability distribution of the distance of attached dynein anchor points from the axis of the IS *r*_IS_ when 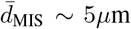. Two dimensional probability density of attached dynein in the IS, 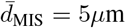. (e) Area density of cortical sliding dyneins 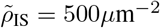. (f) 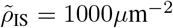.

**Figure 9:**
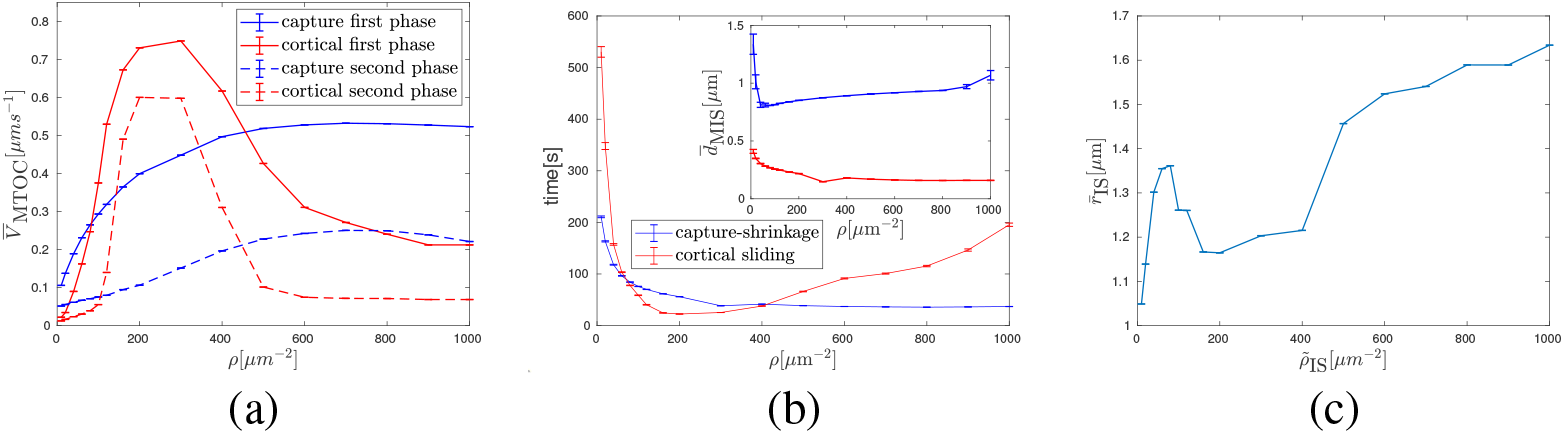
Comparison of the capture-shrinkage and the cortical sliding mechanism in terms of average MTOC velocity in both phases 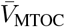, times of repositioning and the final MTOC-IS distance 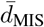. (a) MTOC velocity in the first and the second phase. (b) Repositioning times. Final MTOC-IS distances in the inset. (c) The dependence of average distance 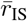 of attached dynein motors from the axis of the IS on dynein area density 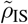 for the case of sole cortical sliding.

Due to the deformation of the cytoskeleton, a large number of MTs is attached to dynein at the end of repositioning, see Fig. 7d and dyneins are predominantly found at the opposite side of the IS(compared to the MTOC). Due to the attachment, the MTOC passes the center of the IS, see Fig. 7a and anchor points of certain dynein motors, see Figs. 6c and 6f. The MTs are attached to anchor points; so, the histogram of *α* changes and the majority of MTs are behind the MTOC, see Fig. 7e. When the MTOC recurs to the IS, the histogram levels, see Fig. 7f and dynein detach.

#### 3.2.3 Cortical sliding with high dynein densities

An example for re-positioning under the effect of cortical sliding with a high dynein density 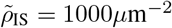 is shown in the supplementary Movie_5. As the area density 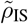 rises, the MTs are more and more attached at the periphery, see Fig. 8d. This is further demonstrated by Figs. 8e and 8f(the center of the IS is almost depopulated when 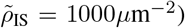. The reason is that there is a sufficient number of dynein to firmly catch MT passing just the periphery of IS. Higher number of MTs also logically means bigger pulling forces on MTs. In a spherical cell, dyneins act in competition which leads to dynein detachment. The bigger the competition is, more frequent is the detachment as can be seen in Fig. 8b, where the highest number of attached dynein corresponds to the lowest area density.

Constantly attaching and detaching dynein does not allow MTs to align with the direction of the MTOC movement. Subsequently, the MTOC “lingers” behind the nucleus before it moves to the IS as the dominant orientation of attached MTs forms slowly. The duration of this inactivity rises with 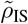, see Fig. 8a. Even when the dominant direction is established, MTs are still attached in every direction slowing down the movement, see Fig. 8c.

Therefore, the slowing in the third section, c.f. Fig. 8a, is caused by two effects. First, the misalignment of MTs resulting in contradictory pulling forces and the lower number of attached dyneins. Second, the increasing probability of catching at the periphery results in MTs being pulled to different places contradicting each other increasingly as the MTOC approaches the IS.

### 3.3 Comparison of cortical sliding and capture-shrinkage

In this section, two mechanisms are compared in terms of the MTOC speeds, times and final the MTOC-IS distances. The biological motivation is that the speed(times) indicates the efficiency of transmission of the force of dynein on the cytoskeleton and the final distance determines the completion of repositioning. In the previous sections, the repositioning was divided into three phases based on the MTOC velocity, see Figs. 3 and 5, which enabled the analysis of the dynamics based on the attached dyneins and deformations of the cytoskeleton structure.

To analyze average speeds, the repositioning is divided into three phases based on the MTOC-IS distance: the activation, the first and the second phase. This approach will later enable a comparison with experimental results. The activation phase ends when *d*_MIS_ < = 8.2*μ*m (identical with the first phase based on the MTOC speed). Although the activation phase is important for the observation of the influence of dynein motors, see Figs. 3c, 5c and 8b, the phase lacks experimental analogy, since in reality the IS alongside with high dynein area density is not created instantly. Therefore, it will not be further analyzed. In the first phase the MTOC-IS distance 8.2*μ*m > *d*_MIS_ > *d*_f_ + 1*μ*m, where *d*_f_ is the final the MTOC-IS distance, which depends on the area density and mechanism. The second phase comprises the last *μ*m of the MTOC journey.

The MTOC speed in the capture-shrinkage repositioning increases with the area density of dyneins for both phases, see Fig. 9a. The development of the average MTOC speed of the cortical sliding repolarization is more difficult since it rises to its maximum(middle densities), see Fig. 3.2.2 and then it falls sharply. The speed of the cortical sliding repositionings is lower except when considering middle area densities of the cortical sliding dynein. Moreover, for the low and high densities, the speed of the capture-shrinkage is more than two times the speed of the cortical sliding, see Fig. 9a. The times of repositionings evolve accordingly, see Fig. 9c. Times are longer for the case of the capture-shrinkage only when *ρ* corresponds to the middle densities of the cortical sliding, see Fig. 9a.

The final MTOC-IS distance decreases with rising *ρ* in the case of the sole capture-shrinkage, see Fig. 9b. In the case of the cortical sliding, the situation is more complicated due to the lack of anchor point. The large final distances at of low area densities are caused by the insufficient pulling force. The shortest distance is at the end of low area densities *ρ* = 80*μ*m^−2^ which is caused by the fact that the formation of the narrow MT stalk, in which MTs pull in alignment, is limited just to low densities, see Figs. 5e, 7c and 8c. Then we can observe a steady rise in final distances caused by growing attachment of MTs at the peripheries as 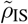 causes increasing competition of pulling forces in the final stages of polarization.

Fig. 9c explains the lower MTOC velocity for cortical sliding. First of all, let us notice that the three regimes of the cortical sliding behavior are visible in 9c. We can see that the increasing 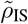 causes MT attachment on periphery of IS, as was already suggested by Figs. 8d 8e and 8f. Moreover, the attached dynein is always predominantly at the periphery, since the average distance for the uniform distribution of dynein is 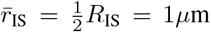. Therefore, as the MTOC approaches the IS, MTs are pulled to different locations, the forces of dynein oppose each other and cause dynein detachment.

The capture-shrinkage mechanism is faster, with the relatively narrow exception of the middle area densities. The cortical sliding never achieves short distances in comparison to the capture-shrinkage; moreover, in the case of high or low densities, the final distances differ substantially. Fig. 9 shows the dependencies on area density. Nevertheless, in the case of the capture-shrinkage, we consider just the density in the center of IS. We should remind that radii of the center and the entire IS are *R*_CIS_ = 0.4*μ*m and *R*_IS_ = 2*μ*m. Since the number of dyneins depends on the area, number of dyneins in the IS *N*_IS_ = 25 * *N*_CIS_, where *N*_CIS_ is the number of dynein in the center of the IS. Consequently, Fig. 9 confirms, that the capture-shrinkage mechanism is the main driving force of the repositioning since this mechanism produces bigger or comparable velocities with just 4% of dynein motors of the cortical sliding. Moreover, the MTOC comes closer to the IS meaning that the capture-shrinkage mechanism is more likely to finish repositioning. To summarize, considering the lower number of dynein, the capture-shrinkage mechanism is largely superior in the considered setup. The most important difference between the two mechanisms is the firm, narrow anchor point in case of the capture-shrinkage mechanism. It assures a firm attachment of the MTs, see Fig. 3f, and a geometrical alignment of the pulling forces in all stages of repositioning. The capture-shrinkage mechanism was identified as the main driving force of repositioning (14) and our model fully support this statement. In the next section we will srutinize the role of cortical sliding.

### 3.4 Combination of Capture-Shrinkage and Cortical Sliding

In this section, the interplay between two mechanisms is analyzed. The comparison of the supplementary movies Movie_6 (capture-shrinkage, with *ρ*_IS_ = 60*μ*m^−2^, 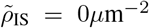) and Movie_7 (both mechanisms combined, with Movie_7, 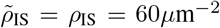) demonstrates the difference between the spindle dynamics under the effect of capture-shrinkage alone and under the effect of both mechanisms combined. They show the first few seconds, just long enough for the MTs to attach to the center of the IS. One clearly sees in Movie_7 that MTs intersecting the IS and attached to cortical sliding dynein, are passed to the center of the IS, where they are captured by cortical sliding dynein. The supplementary Movie_8 shows the complete repositioning of the MTOC under the effect of both mechanism combined 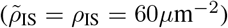

A quantitative analysis in Figs. 10a and 10b shows that the re-polarization speed increases with the cortical-sliding density 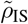 as expected. Quite unexpectedly it turns out that the average number of attached capture-shrinkage dyneins depends on the number of attached cortical sliding dyneins 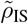: and increases with 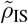, as demonstrated by Fig. 10c.

**Figure 10:**
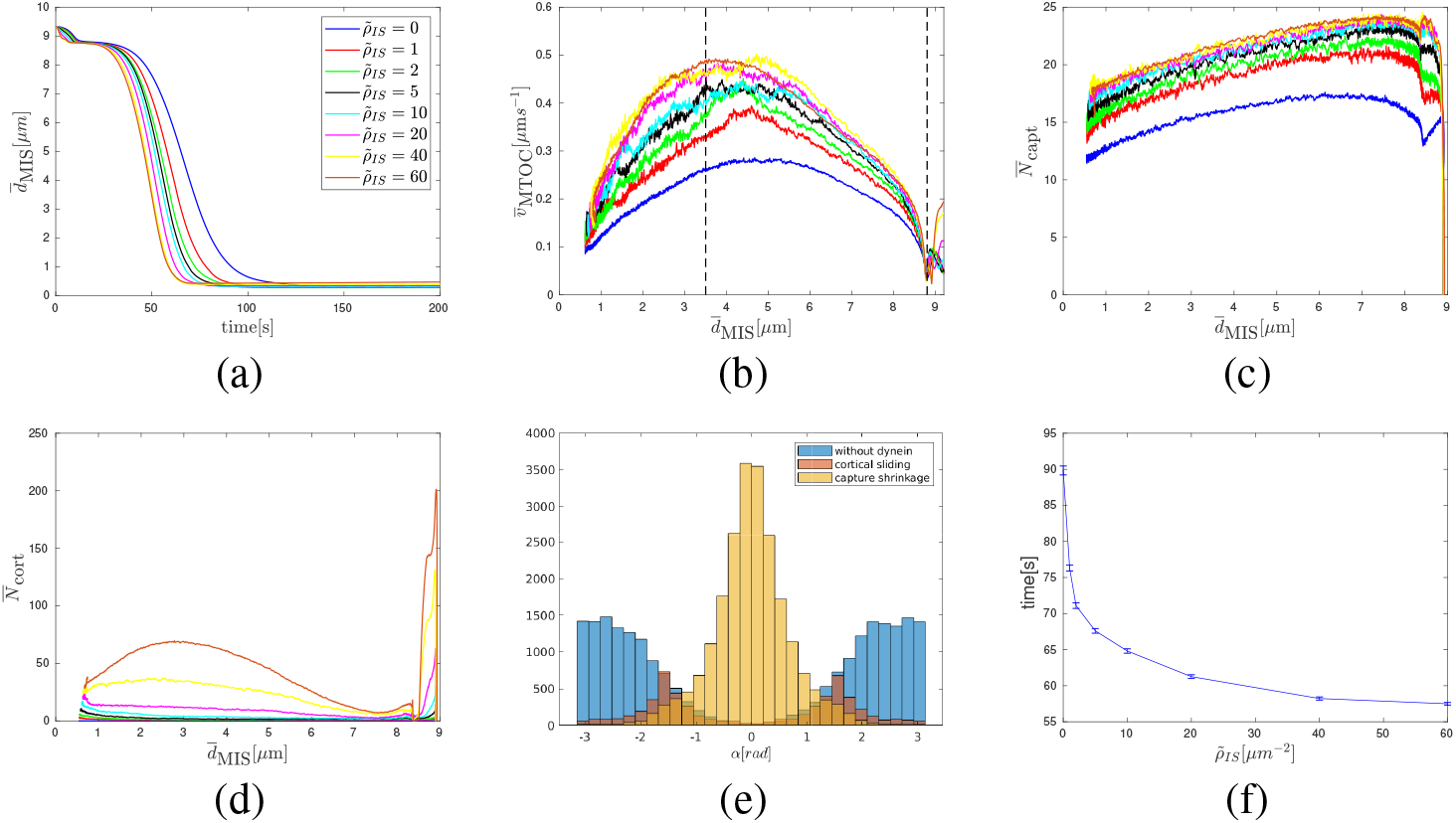
Combination of capture-shrinkage and cortical sliding: (a) Dependence of the average MTOC-IS distance 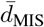 on time. Dependence of average MTOC velocity 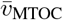 (b), average number of attached capture-shrinkage dyneins 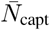 (c) and average number of attached cortical sliding dyneins 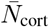 (d) on the average MTOC-IS distance (cortical sliding densities corresponding to different line colors in b-d are the same as in a). (e) Histogram of the angles *α* between the first MT segment and the direction of the MTOC motion. From 500 simulation runs, *t* = 50s, 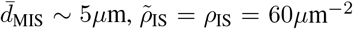. (f) Dependence of times of repositionings on cortical sliding area density 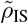.

This finding indicates a synergy of the two mechanisms, capture-shrinkage and cortical sliding, and can be explained by the alignment of the MTs during re-positioning. The MTs attached to the cortical sliding dyneins tend to align with the MTOC movement as demonstrated by the histogram in Fig. 10e, where the the dominant central peak in direction *α* = 0 is caused by capture-shrinkage dyneins and cortical sliding dyneins provide only two small peaks from angles towards the pewriphery of the IS. As MTs align with the MTOC movement, they are captured by the cpature-shrinkage dyneins in the central region of the IS and the number of cortical sliding dyneins drops dramatically as shown in 10d.

A comparison of the histograms shown in 3e for capture-shrinkage alone and 5f for cortical sliding alone reveals the mechanism by which cortical sliding supports capture-shrinkage. Fig. 3f shows that the unattached MTs are pushed back by friction forces, which leads to the opening of the MT spindle, such that MTs cannot intersect the narrow center of IS any more (visualized in Fig. 2f). Attached MTs align with the MTOC movement in the case of the cortical sliding, c.f. Fig. 5f. Therefore, when both mechanisms are combined, MTs attached by the cortical sliding dyneins are not pushed back by friction and they align with the MTOC movement and the attachment probability of capture-shrinkage dynein increases. Comparing the angle histogram for cortical sliding alone, Fig. 5f, with the one for the combined mechanisms, Fig. 10e, demonstrates impressively that most cortical sliding dyneins have detached and attached to capture-shrinkage dyneins.

These observations suggest an answer to the question about the role of cortical sliding: it passes the MTs to the more efficient capture-shrinkage mechanism. Additionally it provides a bigger pulling force than for cortical sliding alone due to the fact that the capture-shrinkage mechanism vice versa also supports the cortical sliding. By comparison of Figs. 5c and 8b with Fig. 10d one realize that the dependencies of the number of cortical sliding dyneins on the MTOC-IS distance are very different. As the MTOC approaches the IS, the number of dyneins acting on MTs decreases in the case of the sole cortical sliding, c.f. Figs. 5c and 8b, but rises for the case of combined mechanisms, c.f. Fig. 10d. The reason lies in the firm anchoring of MTs in the center of the IS and the emergence of the remarkable “arc” formations of attached dynein, c.f. Figs. 11b and 11c.

**Figure 11:**
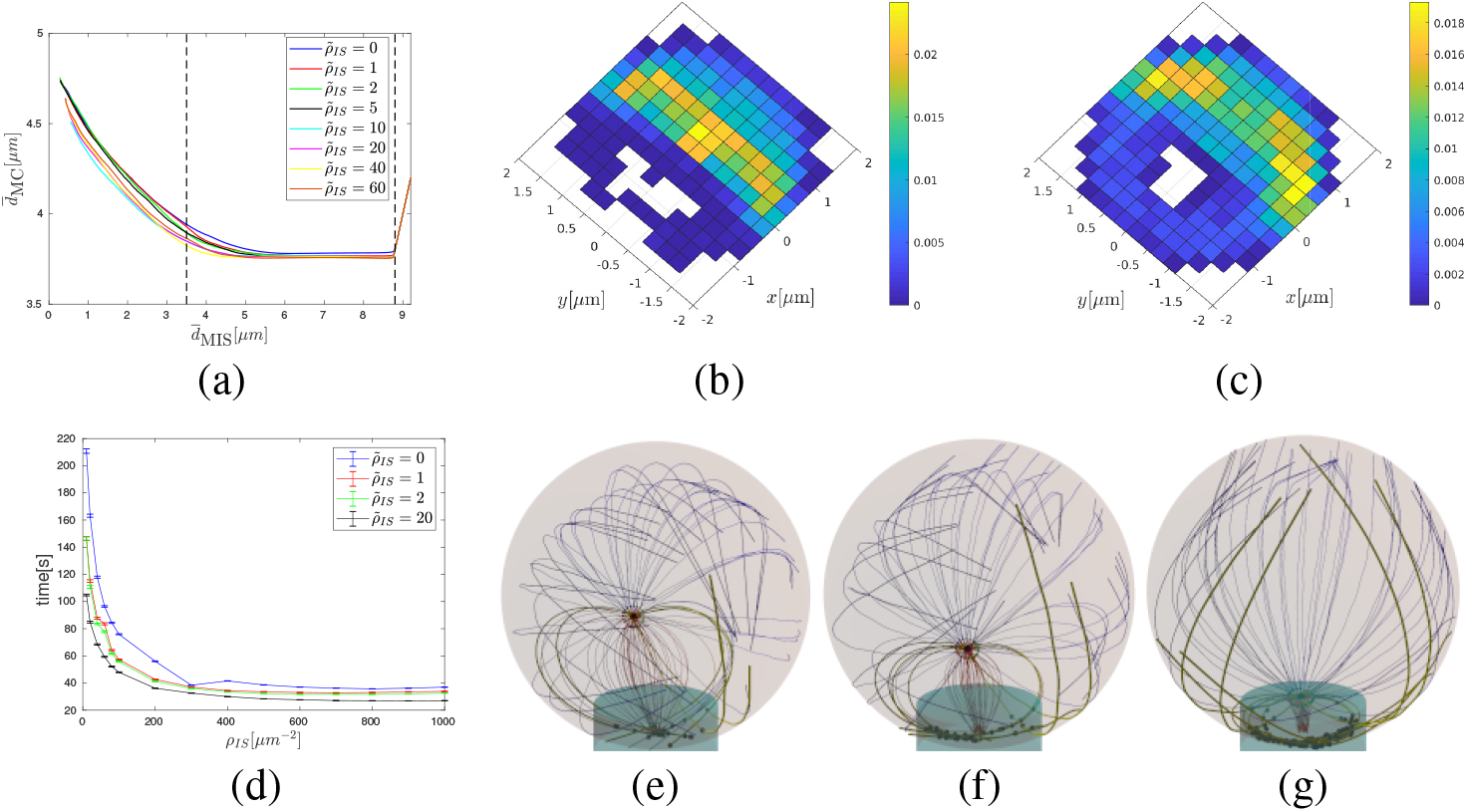
Combination of capture-shrinkage and cortical sliding. (a) Dependence of the average distance between the center of the cell and the MTOC 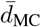 on the average MTOC-IS distance 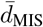. (b)(c) Probability density plot for the spatial distribution of attached dynein. (b) *t* = 50s, 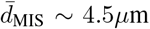. (c) *t* = 60s, 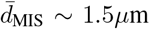. (d) Repositioning times as a function of the density of capture-shrinkage dynein *ρ*_IS_ vfor four different values of the cortical sliding area density *ρ*_IS_. (e),(f),(g) Snapshots from simulation. The blue, red and bold yellow curves correspond to microtubules without dynein, with the capture-shrinkage and the cortical sliding, respectively. Black dots depicts positions of attached dynein motors. (e) *d*_MIS_ = 4.5*μ*m, (f) *d*_MIS_ = 2.5*μ*m,(g) *d*_MIS_ = 1*μ*m

The speed of the capture-shrinkage processes explains this surprising finding. The capture-shrinkage mechanism is more efficient since the MTs shorten due to depolymerization, align with the MTOC movement and they are pulled to the same place. Slower stepping in the cortical sliding mechanism will result in MT lengths between the MTOC and the IS far longer than the direct distance. Therefore, MTs have to bend, see Figs. 11e–11g, which explains the “arc” patterns of attached dynein in the IS. In other words, firm anchoring of capture-shinkage pushes the cortical sliding MTs against the IS, causing further attachment. By comparison of 5d and 11d one observes that the MTOC approaches the IS closer in the case of combined mechanisms than in the case of the cortical sliding, which is another proof of the pulling of the MTOC towards the center of the IS. We conclude that the cortical sliding mechanism supports the dominant capture-shrinkage mechanism by “passing” the MTs, and the capture-shrinkage mechanism supports the cortical sliding mechanism by providing the anchoring and pushing the MTs against the IS.

This synergy is also indicated by Fig. 11d, which shows the total re-positioning times as a function of the density of capture shrinkage dynein for various fixed values of the cortical sliding dynein. Although the re-positioning time does not decrease further for large values of the capture-shrinkage dynein density (*ρ*_IS_ > 600*μm*^−2^) it can actually be decreased further by increasing cortical-sliding dynein. Consequently, the combination of the two mechanisms with relatively low area densities is faster than the dominant mechanism alone with maximum area density (compare cases *ρ*_IS_ = 200*μ*m^−2^ with various 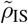 with the case of *ρ*_IS_ = 1000*μ*m^−2^ in Fig. 11d). Further parameter variations supporting this result can be found in the supplementary information 3.3 3. The effect is certainly advantageous for the cell since the cortical sliding mechanism is not as efficient as the capture-shrinkage mechanism and requires a large number of dyneins. considering a large amount of dynein required.

## 4 Discussion

We have analyzed the experiments of (14) with a mechanistic model for the relocation of the MTOC in T-cells. By using biologically realistic values for the model parameters, like the number and stiffness of MTs, dynein pulling forces and detachment probabilities, and cytosol viscosity we can recapitulate for a wide range of dynein densities the experimental observations of (14). In particular the times scale for the completion of the relocation process as well as the MTOC velocities predicted by the model agree well with the experimental results.

Our model predicts that the cytoskeleton deforms substantially during the MTOC-repositioning process due to the combined effects of the capture-shrinkage mechanism and friction forces. The captured MTs form a narrow stalk between the MTOC and the IS, straightening under the tension caused by dynein motors acting on it and causing the rotation of the whole spindle towards it. Concomitantly unattached MTs are pushed backwards by the emerging viscous drag “opening” the spindle, c.f. 2e and 2f. Thus our model provides a mechanistic explanation of the spindle openening that is clearly visible also in the the experiments, as for instance in Fig. 5A of (14). The opening can also be seen in the case of combined mechanisms, although for partially different reasons 11e-11g.

The spindle opening might have interesting consequences for the distribution of Ca^2+^ in the cell, which is highly relevant for cell function. As the cytoskeleton rotates, the mitochondria are dragged with it (16), until they approach IS. Due to spindle opening the mitochondria are positioned asymmetrically around the IS resulting in an asymmetric absorption and redistribution of Ca^2+^ by the mitochondria. Consequently an asymmetric distribution of Ca^2+^ arises around the IS, whose function might deserve further investigation.

The detailed analysis of the spindle arrangement for the cortical sliding mechanism revealed three different deformation characteristics depending on three regimes for the dynein density 3.2. This observation opens the intersting experimental perspective to estimate the dynein distribution from the spindle deformation during MTOC repositioning.

Moreover, our model also predicts a biphasic behavior of the relocation process as reported for the experiments in (14). The Figures 3a and 10a bear a clear resemblance to Figure 3D of (14). We showed that after a short initial period, in which MTs start to attach to dynein, the first phase observed experimentally corresponds in our model to the circular motion around of the MTOC the nucleus and the second phase to to the last 1*μ*m of the MTOC movement, where it detaches from the nucleus and moves more or less straight to the IS with a substantially reduced velocity – for both mechanisms, capture-shrinkage and cortical sliding, and a large range of dynein densities. During the latter phase the MTOC increases its distance from the cell center by approximately 1*μ*m ~ 0.2 * *R*_Cell_, which is close to the value reported in (14).

It was hypothesized in (14) that a resistive force emerges at the transition point between the phases, causing slowing down of the MTOC. Our model shows that the assumption of a resistive force is not necessary to explain the biphasic behavior: the velocity of the MTOC is only determined by the number of motors pulling on the MTs, and on MT alignment 3, 5a and 8. The reason for the slowing down is therefore simply the decrease of the number of dyneins attached to the MTs, which again is a consequence of the changing geometry and forces during the movement of the MTOC, i.e. a consequence of the interplay between spindle and motors.

Experimentally it was also observed that a treatment with taxol substantially reduced the velocity of the repositioning. Taxol impedes depolymerization of the MTs which we could mimic in our model by reducing the capture-shrinkage efficiency. With this modification our model reproduces the experimental observation (SI 10a and 10b).

An interesting prediction of our model is that the two mechanisms, cpature-shrinkage and cortical sliding, appear to act in remarkable synergy 3.4, which provides an answer to the question about the role of the cortical sliding (14). The cortical sliding passes the MTs to the more efficient capture-shrinkage mechanism which in return provides the firm anchor point. Therefore, the cortical sliding is useful even in the configuration, when the capture-shrinkage is dominant. The synergy has a very practical effect since the combination of mechanisms with relatively low area densities can be faster than only the dominant mechanism with much higher area density 11d. Therefore, the synergy of two mechanisms can substantially reduce the area densities necessary for effective repositioning and reduces the necessary resources (dynein). One may speculate that in a real cell obstacles, like organelles that are hard to move, In our model the cytoskeleton does not have to force its way through multiple organelles with complex structure and the synergy manifests itself mainly in the speed of the repositioning process. But one could speculate that in the real cell the synergy could actually can make the difference between completed and no repositioning.

It was proven in (43) that dynein colocalizes with the ADAP ring in the pSMAC. Moreover, in (17) was hypothesized that the MTs are part of the reason why the pSMAC takes the form of the ring. Additionally, (17) reported the sharp turns in MTs upon interaction with the pSMAC and that the MTs do not project directly to the cSMAC. In our model for cortical sliding dynein is homogeneously distributed over the entire IS, nevertheless we observe that dyneins attach to MTs predominantly at the periphery of the IS, c.f. 8d, 8e and 8f. If both mechanisms are present attached cortical sliding dyneins are even completely absent in the central region, c.f. 11. We therefore conclude like (17) that cortical sliding MTs do not project directly into the cSMAC and identify the periphery of the IS as the region where cortical sliding MTs are anchored. In agreement with the experiments (17) we observe that cortical sliding MTs turn upon contact with the periphery 11e-11g, twist and contribute to the formations of dynein “arcs”. Since dynein in the central region of the IS does not contribute to MTOC repositioning via cortical sliding one could hypothesisze that the pSMAC takes the shape of the ring to facilitate interaction with MTs (17).

We presented a numerical analysis of the repositioning in the case when the MTOC and the IS are initially at the opposite sides of the cell. Even the case so restricted brought interesting results enabling the comparison with experiments and proposition of explanation of experimental observable. We found that the cell performs the repositioning with great efficiency. The dyneins are placed only at the peripheries of IS(pSMAC), which is the place where they are used the most, evacuating less used regions. Moreover, we introduced the synergy of two mechanisms that minimizes the necessary area density of dynein.

In this work we presented the results of our theoretical analysis of the MTOC repositioning that is relevant for the experimental setup in (14), where the IS and the intial position of the MTOC are diametrically opposed. Here it turned out that even if both mechanism, capture-shrinkage and cortical sliding, are at work, capture shrinkage is always dominant as reported in (14). In a second part of this work (67), we will examine other initial positions of MTOC and IS that will naturally occur in biologically relevant situations, and we will investigate under which circumstances cortical sliding will become the dominant mechanism over capture-shrinkage. Moreover, we will further demonstrate the synergy of two mechanisms introduced in this work and prove that it has more far-reaching effects in other initial configurations than the one studied here. Also in the situation in which the T cell establishes two IS interesting dynamical behavior of the MTOC can be expected and will be analyzed in detail. In the end, we will see that in T cells the two mechanisms, capture-shrinkage and cortical sliding, and the spatial distribution of dynein are combined such as to minimize the number of dyneins necessary for polarization and to minimize the damage of the MT spindle.

## AUTHOR CONTRIBUTIONS

H.R. designed the research. I.H. performed calculations, prepared figures, and analyzed the data. I.H. and H.R. wrote the manuscript.

## ACKNOWLEDGMENTS

The authors wish to thank Bin Qu and Renping Zhao for helpful discussions.

This work was financially supported by the German Research Foundation (DFG) within the Collaborative Research Center SFB 1027.

## References

1. Rudolph, M. G., R. L. Stanfield, and I. A. Wilson, 2006. How TCRs bind MHCs, peptides, and coreceptors. Annu. Rev. Immunol. 24:419–466.

2. Garcia, K. C., 2012. Reconciling views on T cell receptor germline bias for MHC. Trends Immunol 33:429–436.

3. Zinkernagel, R. M., and P. C. Doherty, 1974. Restriction of in vitro T cell-mediated cytotoxicity in lymphocytic choriomeningitis within a syngeneic or semiallogeneic system. Nature 248:701.

4. Attaf, M., M. Legut, D. K. Cole, and A. K. Sewell, 2015. The T cell antigen receptor: the Swiss army knife of the immune system. Clin Exp Immunol 181:1–18.

5. Wucherpfennig, K. W., 2004. T cell receptor crossreactivity as a general property of T cell recognition. Mol. Immunol. 40:1009–1017.

6. Babbitt, B. P., P. M. Allen, G. Matsueda, E. Haber, and E. R. Unanue, 1985. Binding of immunogenic peptides to Ia histocompatibility molecules. Nature 317:359.

7. Monks, C. R., B. A. Freiberg, H. Kupfer, N. Sciaky, and A. Kupfer, 1998. Three-dimensional segregation of supramolecular activation clusters in T cells. Nature 395:82–86.

8. Dustin, M. L., M. W. Olszowy, A. D. Holdorf, J. Li, S. Bromley, N. Desai, P. Widder, F. Rosenberger, P. A. van der Merwe, P. M. Allen, and A. S. Shaw, 1998. A Novel Adaptor Protein Orchestrates Receptor Patterning and Cytoskeletal Polarity in T-Cell Contacts. Cell 94:667–677.

9. Dustin, M. L., A. K. Chakraborty, and A. S. Shaw, 2010. Understanding the structure and function of the immunological synapse. Cold Spring Harb Perspect Biol 2:a002311.

10. Huang, Y., D. D. Norton, P. Precht, J. L. Martindale, J. K. Burkhardt, and R. L. Wange, 2005. Deficiency of ADAP/Fyb/SLAP-130 destabilizes SKAP55 in Jurkat T cells. J. Biol. Chem. 280:23576–23583.

11. Andre, P., A. M. Benoliel, C. Capo, C. Foa, M. Buferne, C. Boyer, A. M. Schmitt-Verhulst, and P. Bongrand, 1990. Use of conjugates made between a cytolytic T cell clone and target cells to study the redistribution of membrane molecules in cell contact areas. J. Cell Sci. 97:335–347.

12. Geiger, B., D. Rosen, and G. Berke, 1982. Spatial relationships of microtubule-organizing centers and the contact area of cytotoxic T lymphocytes and target cells. J. Cell Biol. 95:137–143.

13. Kupfer, A., D. Louvard, and S. J. Singer, 1982. Polarization of the Golgi apparatus and the microtubule-organizing center in cultured fibroblasts at the edge of an experimental wound. Proc Natl Acad Sci U S A 79:2603–2607.

14. Yi, J., X. Wu, A. H. Chung, J. K. Chen, T. M. Kapoor, and J. A. Hammer, 2013. Centrosome repositioning in T cells is biphasic and driven by microtubule end-on capture-shrinkage. J Cell Biol 202:779–792.

15. Stinchcombe, J. C., E. Majorovits, G. Bossi, S. Fuller, and G. M. Griffiths, 2006. Centrosome polarization delivers secretory granules to the immunological synapse. Nature 443:462–465.

16. Maccari, I., R. Zhao, M. Peglow, K. Schwarz, I. Hornak, M. Pasche, A. Quintana, M. Hoth, B. Qu, and H. Rieger, 2016. Cytoskeleton rotation relocates mitochondria to the immunological synapse and increases calcium signals. Cell Calcium 60:309–321.

17. Kuhn, J. R., and M. Poenie, 2002. Dynamic Polarization of the Microtubule Cytoskeleton during CTL-Mediated Killing. Immunity 16:111–121.

18. Hui, K. L., and A. Upadhyaya, 2017. Dynamic microtubules regulate cellular contractility during T-cell activation. Proc Natl Acad Sci U S A 114:E4175–E4183.

19. Kupfer, A., and G. Dennert, 1984. Reorientation of the microtubule-organizing center and the Golgi apparatus in cloned cytotoxic lymphocytes triggered by binding to lysable target cells. J. Immunol. 133:2762–2766.

20. Kupfer, A., S. L. Swain, C. A. Janeway, and S. J. Singer, 1986. The specific direct interaction of helper T cells and antigen-presenting B cells. Proc Natl Acad Sci U S A 83:6080–6083.

21. Gurel, P. S., A. L. Hatch, and H. N. Higgs, 2014. Connecting the Cytoskeleton to the Endoplasmic Reticulum and Golgi. Curr. Biol. 24:R660–R672.

22. Lee, C., and L. B. Chen, 1988. Dynamic behavior of endoplasmic reticulum in living cells. Cell 54:37–46.

23. Waterman-Storer, C. M., and E. D. Salmon, 1998. Endoplasmic reticulum membrane tubules are distributed by microtubules in living cells using three distinct mechanisms. Curr. Biol. 8:798–806.

24. Palmer, K. J., P. Watson, and D. J. Stephens, 2005. The role of microtubules in transport between the endoplasmic reticulum and Golgi apparatus in mammalian cells. Biochem. Soc. Symp. 1–13.

25. Müllbacher, A., P. Waring, R. T. Hla, T. Tran, S. Chin, T. Stehle, C. Museteanu, and M. M. Simon, 1999. Granzymes are the essential downstream effector molecules for the control of primary virus infections by cytolytic leukocytes. PNAS 96:13950–13955.

26. Lowin, B., M. C. Peitsch, and J. Tschopp, 1995. Perforin and granzymes: crucial effector molecules in cytolytic T lymphocyte and natural killer cell-mediated cytotoxicity. Curr. Top. Microbiol. Immunol. 198:1–24.

27. Voskoboinik, I., M. J. Smyth, and J. A. Trapani, 2006. Perforin-mediated target-cell death and immune homeostasis. Nat. Rev. Immunol. 6:940–952.

28. Grossman, W. J., P. A. Revell, Z. H. Lu, H. Johnson, A. J. Bredemeyer, and T. J. Ley, 2003. The orphan granzymes of humans and mice. Curr. Opin. Immunol. 15:544–552.

29. Krzewski, K., and J. E. Coligan, 2012. Human NK cell lytic granules and regulation of their exocytosis. Front Immunol 3:335.

30. Yannelli, J. R., J. A. Sullivan, G. L. Mandell, and V. H. Engelhard, 1986. Reorientation and fusion of cytotoxic T lymphocyte granules after interaction with target cells as determined by high resolution cinemicrography. J. Immunol. 136:377–382.

31. Pasternack, M. S., C. R. Verret, M. A. Liu, and H. N. Eisen, 1986. Serine esterase in cytolytic T lymphocytes. Nature 322:740.

32. Poo, W. J., L. Conrad, and C. A. Janeway, 1988. Receptor-directed focusing of lymphokine release by helper T cells. Nature 332:378–380.

33. Kupfer, H., C. R. Monks, and A. Kupfer, 1994. Small splenic B cells that bind to antigen-specific T helper (Th) cells and face the site of cytokine production in the Th cells selectively proliferate: immunofluorescence microscopic studies of Th-B antigen-presenting cell interactions. J. Exp. Med. 179:1507–1515.

34. Stinchcombe, J. C., G. Bossi, S. Booth, and G. M. Griffiths, 2001. The immunological synapse of CTL contains a secretory domain and membrane bridges. Immunity 15:751–761.

35. Haddad, E. K., X. Wu, J. A. Hammer, and P. A. Henkart, 2001. Defective granule exocytosis in Rab27a-deficient lymphocytes from Ashen mice. J. Cell Biol. 152:835–842.

36. Griffiths, G. M., 1997. Protein sorting and secretion during CTL killing. Semin. Immunol. 9:109–115.

37. Stinchcombe, J. C., D. C. Barral, E. H. Mules, S. Booth, A. N. Hume, L. M. Machesky, M. C. Seabra, and G. M. Griffiths, 2001. Rab27a is required for regulated secretion in cytotoxic T lymphocytes. J. Cell Biol. 152:825–834.

38. Calvo, V., and M. Izquierdo, 2018. Imaging Polarized Secretory Traffic at the Immune Synapse in Living T Lymphocytes. Front Immunol 9.

39. Kupfer, A., G. Dennert, and S. J. Singer, 1985. The reorientation of the Golgi apparatus and the microtubule-organizing center in the cytotoxic effector cell is a prerequisite in the lysis of bound target cells. J. Mol. Cell. Immunol. 2:37–49.

40. Lin, J., M. J. Miller, and A. S. Shaw, 2005. The c-SMAC. J Cell Biol 170:177–182.

41. Choudhuri, K., and M. L. Dustin, 2010. Signaling microdomains in T cells. FEBS Lett 584.

42. Martín-Cófreces, N. B., J. Robles-Valero, J. R. Cabrero, M. Mittelbrunn, M. Gordón-Alonso, C.-H. Sung, B. Alarcón, J. Vázquez, and F. Sánchez-Madrid, 2008. MTOC translocation modulates IS formation and controls sustained T cell signaling. J. Cell Biol. 182:951–962.

43. Combs, J., S. J. Kim, S. Tan, L. A. Ligon, E. L. F. Holzbaur, J. Kuhn, and M. Poenie, 2006. Recruitment of dynein to the Jurkat immunological synapse. Proc. Natl. Acad. Sci. U.S.A. 103:14883–14888.

44. Hashimoto-Tane, A., T. Yokosuka, K. Sakata-Sogawa, M. Sakuma, C. Ishihara, M. Tokunaga, and T. Saito, 2011. Dynein-Driven Transport of T Cell Receptor Microclusters Regulates Immune Synapse Formation and T Cell Activation. Immunity 34:919–931.

45. Stinchcombe, J. C., and G. M. Griffiths, 2014. Communication, the centrosome and the immunological synapse. Philos Trans R Soc Lond B Biol Sci 369.

46. Quann, E. J., E. Merino, T. Furuta, and M. Huse, 2009. Localized diacylglycerol drives the polarization of the microtubule-organizing center in T cells. Nat. Immunol. 10:627–635.

47. Laan, L., N. Pavin, J. Husson, G. Romet-Lemonne, M. van Duijn, M. P. López, R. D. Vale, F. Jülicher, S. L. Reck-Peterson, and M. Dogterom, 2012. Cortical dynein controls microtubule dynamics to generate pulling forces that position microtubule asters. Cell 148:502–514.

48. Kim, M. J., and I. V. Maly, 2009. Deterministic Mechanical Model of T-Killer Cell Polarization Reproduces the Wandering of Aim between Simultaneously Engaged Targets. PLOS Computational Biology 5:e1000260.

49. Sarkar, A., H. Rieger, and R. Paul, 2019. Search and Capture Efficiency of Dynamic Microtubules for Centrosome Relocation during IS Formation. Biophys. J. 116:2079–2091.

50. Cooper, G. M., and G. M. Cooper, 2000. The Cell. Sinauer Associates, 2nd edition.

51. Li, H., D. J. DeRosier, W. V. Nicholson, E. Nogales, and K. H. Downing, 2002. Microtubule Structure at 8 A Resolution. Structure 10:1317–1328.

52. Meurer-Grob, P., J. Kasparian, and R. H. Wade, 2001. Microtubule Structure at Improved Resolution. Biochemistry 40:8000–8008.

53. Measuring the flexural rigidity of actin filaments and microtubules from their thermal fluctuating shapes: A new perspective | Elsevier Enhanced Reader.

54. Takasone, T., S. Juodkazis, Y. Kawagishi, A. Yamaguchi, S. Matsuo, H. Sakakibara, H. Nakayama, and H. Misawa, 2002. Flexural Rigidity of a Single Microtubule. Jpn. J. Appl. Phys. 41:3015–3019.

55. Gittes, F., B. Mickey, J. Nettleton, and J. Howard, 1993. Flexural rigidity of microtubules and actin filaments measured from thermal fluctuations in shape. J. Cell Biol. 120:923–934.

56. Broedersz, C., and F. MacKintosh, 2014. Modeling semiflexible polymer networks. Rev. Mod. Phys. 86:995–1036.

57. Reconstruction of the Centrosome Cycle from Cryoelectron Micrographs | Elsevier Enhanced Reader.

58. Winey, M., and E. O’Toole, 2014. Centriole structure. Philos Trans R Soc Lond B Biol Sci 369.

59. Guichard, P., V. Hachet, N. Majubu, A. Neves, D. Demurtas, N. Olieric, I. Fluckiger, A. Yamada, K. Kihara, Y. Nishida, S. Moriya, M. O. Steinmetz, Y. Hongoh, and P. Gönczy, 2013. Native architecture of the centriole proximal region reveals features underlying its 9-fold radial symmetry. Curr. Biol. 23:1620–1628.

60. Bernhard, W., and E. De Harven, 1956. [Electron microscopic study of the ultrastructure of centrioles in vertebra]. Z Zellforsch Mikrosk Anat 45:378–398.

61. Woodruff, J. B., O. Wueseke, and A. A. Hyman, 2014. Pericentriolar material structure and dynamics. Philos. Trans. R. Soc. Lond., B, Biol. Sci. 369.

62. Moritz, M., M. B. Braunfeld, V. Guénebaut, J. Heuser, and D. A. Agard, 2000. Structure of the gamma-tubulin ring complex: a template for microtubule nucleation. Nat. Cell Biol. 2:365–370.

63. Robbins, E., G. Jentzsch, and A. Micali, 1968. The Centriole Cycle in Synchronized Hela Cells. J. Cell Biol. 36:329–339.

64. Moritz, M., M. B. Braunfeld, J. W. Sedat, B. Alberts, and D. A. Agard, 1995. Microtubule nucleation by gamma-tubulin-containing rings in the centrosome. Nature 378:638–640.

65. Burgess, S. A., M. L. Walker, H. Sakakibara, P. J. Knight, and K. Oiwa, 2003. Dynein structure and power stroke. Nature 421:715–718.

66. Belyy, V., M. A. Schlager, H. Foster, A. E. Reimer, A. P. Carter, and A. Yildiz, 2016. The mammalian dynein-dynactin complex is a strong opponent to kinesin in a tug-of-war competition. Nat. Cell Biol. 18:1018–1024.

67. H., H. I. R. To be published.

68. Howard, J., 2001. Mechanics of Motor Proteins and the Cytoskeleton. Sinauer Associates is an imprint of Oxford University Press, Sunderland, Mass, new edition edition.

69. Leith, D., 1987. Drag on Nonspherical Objects. Aerosol Science and Technology 6:153–161.

70. Bereiter-Hahn, J., and M. Vöth, 1994. Dynamics of mitochondria in living cells: shape changes, dislocations, fusion, and fission of mitochondria. Microsc. Res. Tech. 27:198–219.

71. Jakobs, S., 2006. High resolution imaging of live mitochondria. Biochimica et Biophysica Acta (BBA) - Molecular Cell Research 1763:561–575.

72. Chaudhuri, A., 2016. Cell Biology by the Numbers. Yale J Biol Med 89:425–426.

73. 1966. An Atlas of Fine Structure. The Cell. Its Organelles and Inclusions. Ann Intern Med 64:968.

74. Jakobs, S., and C. A. Wurm, 2014. Super-resolution microscopy of mitochondria. Current Opinion in Chemical Biology 20:9–15.

75. Xu, H., W. Su, M. Cai, J. Jiang, X. Zeng, and H. Wang, 2013. The Asymmetrical Structure of Golgi Apparatus Membranes Revealed by In situ Atomic Force Microscope. PLoS One 8.

76. Ladinsky, M. S., D. N. Mastronarde, J. R. McIntosh, K. E. Howell, and L. A. Staehelin, 1999. Golgi Structure in Three Dimensions: Functional Insights from the Normal Rat Kidney Cell. J Cell Biol 144:1135–1149.

77. Day, K. J., L. A. Staehelin, and B. S. Glick, 2013. A Three-Stage Model of Golgi Structure and Function. Histochem Cell Biol 140:239–249.

78. Huang, S., and Y. Wang, 2017. Golgi structure formation, function, and post-translational modifications in mammalian cells. F1000Res 6.

79. Westrate, L. M., J. E. Lee, W. A. Prinz, and G. K. Voeltz, 2015. Form follows function: the importance of endoplasmic reticulum shape. Annu. Rev. Biochem. 84:791–811.

80. English, A. R., and G. K. Voeltz, 2013. Endoplasmic reticulum structure and interconnections with other organelles. Cold Spring Harb Perspect Biol 5:a013227.

81. English, A. R., N. Zurek, and G. K. Voeltz, 2009. Peripheral ER structure and function. Curr. Opin. Cell Biol. 21:596–602.

82. Shibata, Y., G. K. Voeltz, and T. A. Rapoport, 2006. Rough sheets and smooth tubules. Cell 126:435–439.

83. Hu, J., W. A. Prinz, and T. A. Rapoport, 2011. Weaving the Web of ER Tubules. Cell 147:1226–1231.

84. Alberts, B., A. Johnson, J. Lewis, M. Raff, K. Roberts, and P. Walter, 2007. Molecular Biology of the Cell, 5th Edition. Garland Science, New York, 5th edition edition.

85. Goodenough, U. W., B. Gebhart, V. Mermall, D. R. Mitchell, and J. E. Heuser, 1987. High-pressure liquid chromatography fractionation of Chlamydomonas dynein extracts and characterization of inner-arm dynein subunits. Journal of Molecular Biology 194:481–494.

86. Gee, M. A., J. E. Heuser, and R. B. Vallee, 1997. An extended microtubule-binding structure within the dynein motor domain. Nature 390:636–639.

87. Goodenough, U., and J. Heuser, 1984. Structural comparison of purified dynein proteins with in situ dynein arms. Journal of Molecular Biology 180:1083–1118.

88. Schmidt, H., E. S. Gleave, and A. P. Carter, 2012. Insights into dynein motor domain function from a 3.3 A crystal structure. Nat Struct Mol Biol 19:492–S1.

89. Leduc, C., O. Campàs, K. B. Zeldovich, A. Roux, P. Jolimaitre, L. Bourel-Bonnet, B. Goud, J.-F. Joanny, P. Bassereau, and J. Prost, 2004. Cooperative extraction of membrane nanotubes by molecular motors. Proc. Natl. Acad. Sci. U.S.A. 101:17096–17101.

90. Kamiya, N., T. Mashimo, Y. Takano, T. Kon, G. Kurisu, and H. Nakamura, 2016. Elastic properties of dynein motor domain obtained from all-atom molecular dynamics simulations. Protein Eng Des Sel 29:317–325.

91. Lindemann, C. B., and A. J. Hunt, 2003. Does axonemal dynein push, pull, or oscillate? Cell Motil. Cytoskeleton 56:237–244.

92. Sakakibara, H., H. Kojima, Y. Sakai, E. Katayama, and K. Oiwa, 1999. Inner-arm dynein c of Chlamydomonas flagella is a single-headed processive motor. Nature 400:586–590.

93. Sakakibara, H., and K. Oiwa, 2011. Molecular organization and force-generating mechanism of dynein. The FEBS Journal 278:2964–2979.

94. Gennerich, A., A. P. Carter, S. L. Reck-Peterson, and R. D. Vale, 2007. Force-Induced Bidirectional Stepping of Cytoplasmic Dynein. Cell 131:952–965.

95. Toba, S., T. M. Watanabe, L. Yamaguchi-Okimoto, Y. Y. Toyoshima, and H. Higuchi, 2006. Overlapping hand-overhand mechanism of single molecular motility of cytoplasmic dynein. PNAS 103:5741–5745.

96. Mallik, R., D. Petrov, S. A. Lex, S. J. King, and S. P. Gross, 2005. Building complexity: an in vitro study of cytoplasmic dynein with in vivo implications. Curr. Biol. 15:2075–2085.

97. Mallik, R., B. C. Carter, S. A. Lex, S. J. King, and S. P. Gross, 2004. Cytoplasmic dynein functions as a gear in response to load. Nature 427:649–652.

98. Reck-Peterson, S. L., A. Yildiz, A. P. Carter, A. Gennerich, N. Zhang, and R. D. Vale, 2006. Single-Molecule Analysis of Dynein Processivity and Stepping Behavior. Cell 126:335–348.

99. Kural, C., H. Kim, S. Syed, G. Goshima, V. I. Gelfand, and P. R. Selvin, 2005. Kinesin and dynein move a peroxisome in vivo: a tug-of-war or coordinated movement? Science 308:1469–1472.

100. Torisawa, T., M. Ichikawa, A. Furuta, K. Saito, K. Oiwa, H. Kojima, Y. Y. Toyoshima, and K. Furuta, 2014. Autoinhibition and cooperative activation mechanisms of cytoplasmic dynein. Nature Cell Biology 16:1118–1124.

101. Müller, M. J. I., S. Klumpp, and R. Lipowsky, 2008. Tug-of-war as a cooperative mechanism for bidirectional cargo transport by molecular motors. Proc. Natl. Acad. Sci. U.S.A. 105:4609–4614.

102. King, S. J., and T. A. Schroer, 2000. Dynactin increases the processivity of the cytoplasmic dynein motor. Nat Cell Biol 2:20–24.

103. Nishiura, M., T. Kon, K. Shiroguchi, R. Ohkura, T. Shima, Y. Y. Toyoshima, and K. Sutoh, 2004. A single-headed recombinant fragment of Dictyostelium cytoplasmic dynein can drive the robust sliding of microtubules. J. Biol. Chem. 279:22799–22802.

104. Kon, T., M. Nishiura, R. Ohkura, Y. Y. Toyoshima, and K. Sutoh, 2004. Distinct Functions of Nucleotide-Binding/Hydrolysis Sites in the Four AAA Modules of Cytoplasmic Dynein. Biochemistry 43:11266–11274.

105. Cho, C., S. L. Reck-Peterson, and R. D. Vale, 2008. Regulatory ATPase sites of cytoplasmic dynein affect processivity and force generation. J. Biol. Chem. 283:25839–25845.

106. Kikushima, K., T. Yagi, and R. Kamiya, 2004. Slow ADP-dependent acceleration of microtubule translocation produced by an axonemal dynein. FEBS Letters 563:119–122.

107. Walter, W. J., B. Brenner, and W. Steffen, 2010. Cytoplasmic dynein is not a conventional processive motor. J. Struct. Biol. 170:266–269.

108. Ikuta, J., N. K. Kamisetty, H. Shintaku, H. Kotera, T. Kon, and R. Yokokawa, 2014. Tug-of-war of microtubule filaments at the boundary of a kinesin- and dynein-patterned surface. Scientific Reports 4:5281.

109. Kunwar, A., S. K. Tripathy, J. Xu, M. K. Mattson, P. Anand, R. Sigua, M. Vershinin, R. J. McKenney, C. C. Yu, A. Mogilner, and S. P. Gross, 2011. Mechanical stochastic tug-of-war models cannot explain bidirectional lipid-droplet transport. Proc. Natl. Acad. Sci. U.S.A. 108:18960–18965.

110. Klein, S., C. Appert-Rolland, and L. Santen, 2015. Motility states in bidirectional cargo transport. EPL 111:68005.

111. Montesi, A., D. C. Morse, and M. Pasquali, 2005. Brownian dynamics algorithm for bead-rod semiflexible chain with anisotropic friction. J. Chem. Phys. 122:084903.

112. Fixman, M., 1978. Simulation of polymer dynamics. I. General theory. J. Chem. Phys. 69:1527–1537.

113. Hinch, E. J., 1994. Brownian motion with stiff bonds and rigid constraints. Journal of Fluid Mechanics 271:219–234.

114. Grassia, P. S., E. J. Hinch, and L. C. Nitsche, 1995. Computer simulations of Brownian motion of complex systems. Journal of Fluid Mechanics 282:373–403.

115. Grassia, P., and E. J. Hinch, 1996. Computer simulations of polymer chain relaxation via Brownian motion. Journal of Fluid Mechanics 308:255–288.

116. Pasquali, M., and D. C. Morse, 2002. An efficient algorithm for metric correction forces in simulations of linear polymers with constrained bond lengths. The Journal of Chemical Physics 116:1834–1838.

117. Nedelec, F., and D. Foethke, 2007. Collective Langevin dynamics of flexible cytoskeletal fibers. New J. Phys. 9:427–427.

