## Supplementary material for "Stochastic model of T Cell repolarization during target elimination (I)"

### Stochastic model of T Cell repolarization during target elimination (I), SUPPLEMENTARY MATERIAL

November 6, 2019

### 1 Model of the cell

#### 1.1 Microtubules

The microtubules(MTs) are represented as semiflexible filaments, therefore the Hamiltonian of a single MT is given by:

$$H = \frac{\kappa}{2}\int_0^L \left|\frac{\partial \vec{t}}{\partial s}\right|^2, \tag{1}$$

where $\kappa$ is bending rigidity( $2.2 * 10^{-23}\text{Nm}^2$ ), $L$ is the length of MT, $s$ is arc length, $\vec{t} = \frac{\partial \vec{r}}{\partial s}$ is unit tangent vector and $\vec{r}(s)$ is a position (1). A single MT is represented as a chain of $N$ beads with coordinates $\vec{r}_1, ..., \vec{r}_N$ connected by $N-1$ tangents $\vec{ti} = \vec{r}_{i+1} - \vec{r}_i$ of the length $k = L/(N-1)$. In the present model, the length $k = 0.8\mu$m was used. Since the MT is an inextensible polymer discretized into $N$ beads, $N-1$ constraints must be fulfilled:

$$C_i^{\text{micro}} = |\vec{r}_i - \vec{r}_{i+1}| = k \qquad\qquad i = 1, ..., N-1 \tag{2}$$

The length of the MTs varies considerably, but since short MTs are not relevant for repositioning, we include only just MTs that reach from the MTOC to the IS in the first seconds of repositioning. The maximum length of a MT should be $L > \pi * R_{\text{Cell}}$ to always reach from MTOC to IS. In order to reach the IS in the first stages of repolarization, the length must be $L > \frac{3}{4} * \pi * R_{\text{Cell}}$. Thus, the number of beads $N$ is uniformly distributed between 15 and 20.

##### 1.1.1 Bending forces of the microtubule

The Hamiltonian for the discretized MT can be expressed as:

$$H_{bend} = \kappa_d \sum_{i=0}^{N-2}\left(1 - \frac{\vec{t}_i\vec{t}_{i+1}}{|\vec{t}_i||\vec{t}_{i+1}|}\right), \tag{3}$$

where $\kappa_d = \kappa/k$ is the bending rigidity of the discretized model. The bending force acting on bead $i$ is the derivative of the discretized Hamiltonian with the respect to $\vec{r}_i$:

$$\vec{F}_i^{\text{bend}} = -\frac{\partial H_{bend}}{\partial \vec{r}_i}. \tag{4}$$

If we consider the simplest case (sketched in Figure 2) of three points with coordinates $\vec{r}_1$, $\vec{r}_2$ and $\vec{r}_3$ connected with the tangents $\vec{t}_1$ and $\vec{t}_2$, the bending forces acting on the beads are:

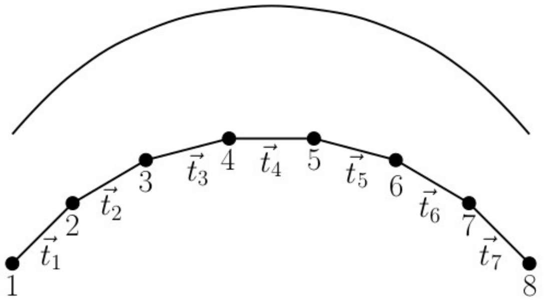


(a)

Figure 1: Bead-rod model of microtubule. The microtubule is divided into 8 point connected by 7 rods. The circles correspond to the positions of the beads. The lengths of rods connecting beads remain constant.

$$\vec{F}_1^{\text{bend}} = \frac{\kappa_d}{|\vec{t}_1|}\left(-\frac{\vec{t}_2}{|\vec{t}_2|} + \frac{\vec{t}_1}{|\vec{t}_1|}\left(\frac{\vec{t}_1\cdot\vec{t}_2}{|\vec{t}_2||\vec{t}_1|}\right)\right), \tag{5a}$$

$$\vec{F}_3^{\text{bend}} = \frac{\kappa_d}{|\vec{t}_2|}\left(\frac{\vec{t}_1}{|\vec{t}_1|} - \frac{\vec{t}_2}{|\vec{t}_2|}\left(\frac{\vec{t}_1\cdot\vec{t}_2}{|\vec{t}_2||\vec{t}_1|}\right)\right) \tag{5b}$$

$$\vec{F}_2^{\text{bend}} = -\vec{F}_3^{\text{bend}} - \vec{F}_1^{\text{bend}}. \tag{5c}$$

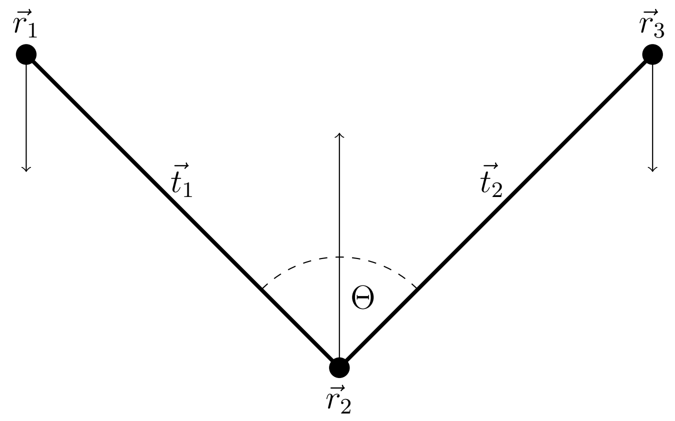


Figure 2: The sketch of bending forces acting on three beads. The filled circles are the microtubule beads. The bending force is determined by the angle $\Theta$ between the tangents $\vec{t}_1$ and $\vec{t}_2$. The narrow arows depict bending forces acting on beads.

###### 1.1.2 Drag coefficient

The drag force on an object depends on the speed of the object, the viscosity $\mu$ of the fluid, the size, and the shape of the object. The MT is divided into segments of a cylindrical shape whose length is substantially bigger than the diameter. The drag depends on the object's area projected normal to the direction of the motion. Therefore, drag forces for the motions parallel and perpendicular to the object are defined as $F = \gamma_{||}\dot{x}$ and $F = \gamma_{\perp}\dot{x}$, respectively, where $\dot{x}$ is speed of the object. Parallel $\gamma_{||}$ and perpendicular $\gamma_{\perp}$ drag coefficients (2) are defined for the case of cylinder in the fluid with viscosity $\mu$ as:

$$\gamma_{||} = \frac{2\pi\mu k}{\ln(k/d) - 0.2} \tag{6a}$$

$$\gamma_{\perp} = \frac{4\pi\mu k}{\ln(k/d) + 0.84}, \tag{6b}$$

where $k$ and $d$ are the length and the diameter of the cylinder, respectively. Therefore, the friction of the cylinder is anisotropic $\gamma_{\perp} \sim 2\gamma_{||}$. However, the anisotropy is hard to implement, as the orientation of segments varies. Therefore, the anisotropy is not implemented. For the case of simplicity, the perpendicular drag coefficient is considered in the simulation to the beads of MT and the same drag coefficient is attributed to all MTOC points.

###### 1.1.3 Drag force of organelles

EM, Golgi and mitochondria are the organelles with different structures and sizes. However, their drag force can be estimated from their volume and surfaces. A dynamic shape factor(3), $K_{\text{sf}}$, can be defined to calculate the drag coefficient of the nonspherical particle of any shape:

$$\gamma = 3\pi d_v \mu K_{\text{sf}}, \tag{7}$$

where $d_v$ is the diameter of the sphere with the same volume as the object. The drag force can be divided between the form drag, coming from the pressure on the surface, and tangential shear stress. Form drag is determined by the objects area projected normal to the direction of the motion. It can be expressed through the Stokes law form drag on a sphere, whose projected area equals to the projected area of a nonspherical object. The diameter of such a sphere is $d_n$. The friction force on the surface can be expressed by the friction on the sphere with the same effective surface. The diameter of the cell is $d_s$. The dynamic shape factor can be defined as:

$$K_{\text{sf}} = \frac{d_n + 2d_s}{3d_v}. \tag{8}$$

The number of mitochondria were measured(just the case of one cell) in (4) ( on average 44 mitochondria in a T-Cell). The size and shape of MTs varies greatly, since they can shrink, grow, go through fission and fusion (5–9). We consider the most common spherocylindrical shape approximated by the cylinder, whose diameter and the length were estimated as $0.75\mu$m and $1.5\mu$m, respectively. Golgi apparatus is very complex structure composed from multiple classes of cisternae differing in form, function and composition that are stacked in various ways (10–13). The endoplasmic reticulum (ER) is also a complex organelle composed from a bilayer forming nuclear envelope and network of sheets and dynamic tubules (14–18). Golgi, ER and mitochondria are connected to cytoskeleton (19) and (4).

The sizes and shapes of EM and Golgi in T-Cell were not measured. For the rough approximate evaluation of drag force we consider the estimates from (20), table 12. The major organelles including EM and Golgi are close to the center and they are in contact. Consequently, effective viscosity in the regions close to the nucleus of the cell is different from more aqueous domain close to the cell membrane where the rotation of the cytoskeleton takes place. Therefore, we assume that the effective viscosity of the medium in which EM and Golgi travel is $\mu_2 = 10 * \mu$. We will express the drag coefficient of organelles as a function of viscosity and compare it with the drag coefficient of the cytoskeleton. For the estimate of the cytoskeleton drag coefficient, the cytoskeleton of 100 MTs was considered and the drag coefficient were calculated using (8).

$$\gamma_{\text{GSER}} \sim 0.00131\mu \tag{9}$$
$$\gamma_{\text{RER}} \sim 0.00160\mu \tag{10}$$
$$\gamma_{\text{Mito}} \sim 0.00257\mu \tag{11}$$
$$\gamma_{\text{Cyto}} \sim 0.00270\mu \tag{12}$$

From the equations (9) it can be seen that the $\gamma_{\text{total}} = (\gamma_{\text{Cyto}} + \gamma_{\text{Mito}} + \gamma_{\text{RER}} + \gamma_{\text{GSER}}) \sim 3 * \gamma_{\text{Cyto}}$. Consequently, to consider the drag force from the organelles in the cell, the drag coefficient of MT is tripled. Thus, the equation (6) can be rewritten as:

$$\gamma_{\parallel} = 3 * \frac{2\pi\mu k}{\ln(k/d) - 0.2} \tag{13a}$$
$$\gamma_{\perp} = 3 * \frac{4\pi\mu k}{\ln(k/d) + 0.84}. \tag{13b}$$

##### 1.1.4 Confinement of the cytoskeleton

The cytoskeleton can move between the wall of the cell and the nucleus. They have a spherical shape and they are modeled as force fields. The force of the wall is null if $|\vec{r}_i| <= R$. Otherwise, the force acting on the segment of the MT or the MTOC is calculated following the subsequent equation:

$$\vec{F}_i^{\text{wall}} = -1\frac{\vec{r}_i}{|\vec{r}_i|}k_1\exp(k_2(|\vec{r}_i| - R)), \tag{14}$$

where $k_1 = 20\text{pN}\mu\text{m}^{-1}$ and $k_2 = 1\text{m}^{-1}$ are chosen constants. The force of nucleus is null if $|\vec{r}_i| > R_{nucleus}$(radius of the nucleus). Otherwise, it can be expressed by:

$$\vec{F}_i^{\text{nucleus}} = \frac{\vec{r}_i}{|\vec{r}_i|}k_1\exp(k_2(R_{nucleus} - |\vec{r}_i|)). \tag{15}$$

##### 1.1.5 Dynein motors

Unfortunately, since the results from the measurements differ greatly, the mechanical properties of dynein remain uncertain. Therefore, the parameters in this section are estimations. The dynein has an anchor and attachment points connected by a stalk. The anchor point has a stable position and the attachment point walks on the MT. The force acting on the MT is determined by the length of the stalk, whose relaxed length was estimates as $L_0 = 15n$m (21–24). Unattached dynein is represented just with one point on the surface of the cell. If the dynein is closer to the MT than $L_0$, the motor protein can attach to the filament. Fluctuations of the membrane can move the dynein motor to the MT. Therefore, attachment probability is defined as:

$$p_a = 5\text{s}^{-1} \qquad d_{md} <= L_0 \tag{16}$$
$$p_a = 5 \cdot \exp((d_{md} - L_0)/p_d)\text{s}^{-1} \qquad d_{md} > L_0, \tag{17}$$

where $d_{md}$ is the distance of the dynein point to the closest point of the MT and $p_d = 10^{-7}$ is a chosen parameter. If the MT is attached, the anchor and attachment points of the dynein motor are placed to the same point on the MT. Attachment probability of dynein is unknown; therefore, the attachment probability $p_a$ corresponding to the attachment ratio of kinesin is considered (25). The force of dynein motor comes from the elastic properties of the stalk:

$$|F_i^{\text{Dynein}}| = 0, \qquad |\vec{r}_{\text{Dynein}}| < L_0 \tag{18}$$
$$\vec{F}_i^{\text{Dynein}} = k_{\text{Dynein}}(|\vec{r}_{\text{Dynein}}| - L_0)\frac{\vec{r}_{\text{Dynein}}}{|\vec{r}_{\text{Dynein}}|} \qquad |\vec{r}_{\text{Dynein}}| > L_0, \tag{19}$$

where $\vec{r}_{\text{Dynein}} = \vec{r}_{\text{anchor}} - \vec{r}_{\text{attach}}$ is the distance between the anchor and the attachment points. The measurements of elastic modulus of the stalk differ (26–29) , therefore, the elastic modulus was estimated to $k_{\text{Dynein}} = 400p\text{N}\mu\text{m}^{-1}$ (30). If the force is null or parallel to the preferred direction of stepping, the probability of stepping to the minus end is:

$$p_- = \frac{V_F}{d_{\text{step}}}, \tag{20}$$

where $V_F$ is the forward speed of dynein and $d_{\text{step}}$ is the length of the step. The steps of dynein have multiple lengths (31–36). Nevertheless, just the most frequently measured length $d_{\text{step}} = 8\text{nm}$ is considered. The forward speed was estimated or measured in various sources (31, 32, 37–43). For our purposes it is estimated to be $V_F = 1000\text{nms}^{-1}$. In the case of the force of the dynein being in the opposite direction to the preferred movement and smaller than a stall force $F_S$, the attachment point steps to the minus end with probability:

$$p_- = \frac{V_F}{d_{\text{step}}}(1 - \frac{|F^{\text{Dynein}}|}{F_S}). \tag{21}$$

The value of the stall force varies greatly (31, 31–34, 38, 44) and (45), we estimate it as $F_S = 4\text{pN}$. If the force aims to the plus end and it is bigger than the stall force, dynein steps to the plus end with probability:

$$p_+ = \frac{V_B}{d_{\text{step}}}. \tag{22}$$

Backward stepping speed is force dependent (31) and the measured values also differ (38), (46). Our estimate is $V_B = 6.0\text{nm}\cdot\text{s}^{-1}$ (31).
The probability of detachment is expressed as:

$$p_{\text{detach}} = \exp(\frac{|F_d|}{F_D}), \tag{23}$$

where the detachment force (38, 46, 47) was estimated as $F_D = F_S/2$. The detached motor is projected to the membrane of the cell. When the attachment point of dynein motor is not on a bead of MT, the force is acting on a point of a segment between two beads. In such a case, the force has to be transmitted to the two closest beads. Since the mechanism of stepping and detachment of dynein is uncertain, we use the model for kinesin stepping (48).

Dynein plays a role in two mechanisms. During cortical sliding mechanism acting in the whole IS, the MT slides on the membrane and its plus-end remains free. MT depolimerizes in the fixed position on the membrane of the cell during capture-shrinkage mechanism acting in the center of the IS. Without the effects of dynein, MT detaches from fixed position.

#### 1.2 Microtubule organizing center (MTOC)

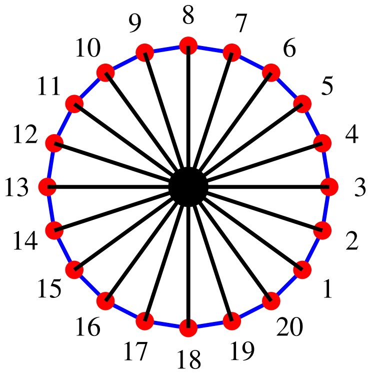


Figure 3: MTOC with $N_{\text{MTOC}} = 20$ MT sprouting points are represented by red spheres. The black sphere in the middle is the center of the MTOC. Black lines connecting the points with the center and blue lines connecting the neighboring points are non-deformable.

The MTOC is modeled as a planar, polygonal structure, composed from so-called sprouting points(points of MT sprouting). If MTOC has $Q^{\text{MTOC}}$ sprouting points, then the equal number of constraints holds the sprouting points in a specified distance from MTOC center(black lines in Figure 3). Therefore, the $i$th constraint is defined as:

$$C_i^{\text{MTOC}} = |\vec{r}_i^{\text{mtoc}} - \vec{r}_c| = R_{\text{MTOC}} \qquad i = 1, ..., Q^{\text{MTOC}}, \tag{24}$$

where $\vec{r}_i^{\text{mtoc}}$ is the position of $i$th sprouting point and $\vec{r}_c$ is the position of the center of the MTOC. Moreover, additional $Q^{\text{MTOC}}$ bonds keep the neighboring points in a constant distance $d^{\text{MTOC}}$(blue lines in Figure 3).

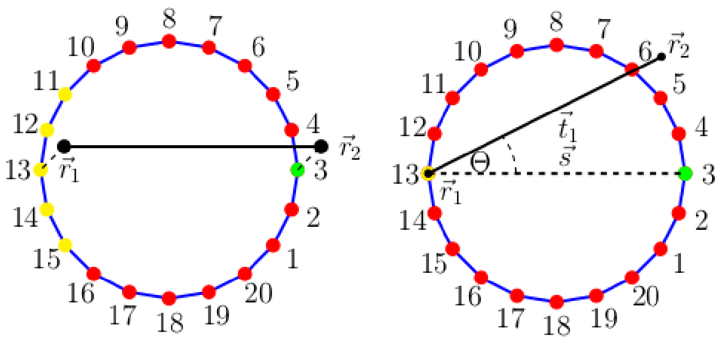


(b) (c)

Figure 4: The scheme of the MTOC composed of 20 sprouting points. The black line denotes the first segment of microtubule. (a) Dashed lines depict the elastic forces connecting two points of microtubule to two MTOC points. The green point represents the sprouting point and yellow points depict possible rear points. (b) Bending force minimizes the angle $\Theta$ between the first segment of microtubule $\vec{t}_1$ and the line $\vec{s}$ connecting sprouting(green) and rear(yellow) point.

$$C_i^{\text{MTOC}} = |\vec{r}_i^{\,\text{mtoc}} - \vec{r}_{i+1}^{\,\text{mtoc}}| = d^{\text{MTOC}} \qquad i = Q^{\text{MTOC}} + 1, ..., 2 \cdot Q^{\text{MTOC}}, \tag{25}$$

When MT is created, the so-called "sprouting point" and "rear point" on the MTOC are chosen 4. The second bead of the MT is attached to the sprouting point and the first bead to the rear point. Consequently, the original MT orientation is given by the direction from the rear point to the sprouting point. Every MTOC point is the sprouting point to the same number of MTs. The rear point is chosen from the points at the approximately opposite side(relatively to sprouting point) of the MTOC 4a, which gives a variety of the initial MT orientations. Elastic force 4a anchors the MT in the MTOC, while bending force 4b forces the MT to be aligned with the line connecting sprouting and rear points. The combination of two forces assures anchoring of the MT and limitation of changes in orientation, simulating the effect of PCM.

##### 1.2.1 Connecting microtubule and MTOC

The first segment of the MT is inside the MTOC 4a. The second and the first the MT beads are attached by elastic forces to the sprouting point and the rear point, respectively 4a. Elastic forces acting on a MT bead can be written as:

$$\vec{F}_2^{\text{elas}} = k_3 |\vec{d}_2| \cdot \frac{\vec{d}_2}{|\vec{d}_2|}, \tag{26}$$

where $k_3 = 1 \cdot \text{pN}\mu\text{m}^{-1}$ is a spring constant and $\vec{d}_2 = \vec{r}_s^{\,\text{MTOC}} - \vec{r}_2^{\,\text{micro}}$, where $\vec{r}_s^{\,\text{MTOC}}$ and $\vec{r}_1^{\,\text{micro}}$ are MTOC sprouting point and the second bead of MT, respectively. Analogically, we can define the forces between the first bead of the MT and the MTOC rear point. Bending forces 4b are calculated via:

$$\vec{F}_{\text{MTOC}}^{\text{bend}} = \frac{\kappa}{|\vec{s}|^2} \left( -\frac{\vec{t}_1}{|\vec{t}_1|} + \frac{\vec{s}}{|\vec{s}|} \left( \frac{\vec{t}_1 \cdot \vec{s}}{|\vec{s}||\vec{t}_1|} \right) \right) \tag{27a}$$

$$\vec{F}_{\text{micro}}^{\text{bend}} = \frac{\kappa}{|\vec{t}_1|^2} \left( \frac{\vec{s}}{|\vec{s}|} - \frac{\vec{t}_1}{|\vec{t}_1|} \left( \frac{\vec{t}_1 \cdot \vec{s}}{|\vec{s}||\vec{t}_1|} \right) \right), \tag{27b}$$

$$\vec{F}_0 = -\vec{F}_{\text{micro}}^{\text{bend}} - \vec{F}_{\text{MTOC}}^{\text{bend}} \tag{27c}$$

where $\vec{s}$ is the segment between the two beads of the MTOC, $\vec{t}_1$ is the first segment of MT. The forces $\vec{F}_{\text{MTOC}}^{\text{bend}}$ and $\vec{F}_{\text{micro}}^{\text{bend}}$ act on the sprouting point and the second bead of MT, respectively. The force $\vec{F}_0$ acts on the first bead of the MT and the rear point.

#### 2 Constrained Langevin dynamics

Using Langevin dynamics, the motion of an unconstrained particle with the position $x_i$ can be expressed:

$$\gamma_i \dot{x}_i = f_i + \eta_i, \tag{28}$$

where $\eta_i$ is a random Langevin force, which is a stochastic, non-differentiable function of time that integrates random interactions with the molecules of the solvent. The force $f_i$ is the sum of all other forces and it depends on the object. The drag coefficient $\gamma$, forces $\eta_i$ and $f_i$ will be defined later for the case of MT and MTOC. In a constrained case, N beads in 3D have to satisfy $Q$ constraints (49):

$$C_a(x_1, ..., x_{3*N}) = c_a \qquad a = 1, ...., Q \tag{29}$$

The constraints have to remain constant in every instance. Therefore, the movement of the beads must satisfy:

$$0 = \dot{C}_a = n_{ia} \cdot \dot{x}_i \qquad a = 1, ...., Q \tag{30}$$

where

$$n_{ia} = \frac{\partial C_a}{\partial x_i}. \tag{31}$$

The motion of constrained bead can be expressed:

$$\gamma \dot{x}_i = f_i + \eta_i - n_{ia}\lambda_a, \tag{32}$$

where $\lambda_a$ is a constraint force conjugate to constraint $\mu$.

#### 2.1 Mid-Step algorithm

Mid-Step algorithm was proposed by Fixman and further generalized by Hinch and Grassia (50–53) The algorithms was elaborated for specific cases by Morse and Pasquali (49) and (54). Using the mobility tensor

$$H_{ik}\gamma = \mathbf{I}_{ik}, \tag{33}$$

the equation (32) can be rewritten as:

$$\dot{x}_i = H_{ij}[F^u_j - n_{ja}\lambda_a], \tag{34}$$

where $F^u_j = f_i + \eta_i$ is unconstrained force. The values of $\lambda_a$ for $a = 1, ..., Q$ can be calculated from the conditions (30) at every instant. It will result in the set of algebraic equations:

$$G_{a\nu}\lambda_\nu = n_{ia}H_{ij}F^u_j, \tag{35}$$

where

$$G_{a\nu} = n_{ia}H_{ij}n_{j\nu}. \tag{36}$$

If the constraint forces are expressed by (35), we get the equation of motion from (32):

$$\dot{x}_i = P_{ij}H_{jk}F^u_k, \tag{37}$$

where

$$P_{ij} = \mathbf{I}_{ij} - H_{ik}n_{ka}G^{-1}_{a\nu}n_{j\nu} \tag{38}$$

is a projection operator. In the case when the mobility tensor is expressed by (33), equation 38 can be rewritten as:

$$P_{ij} = \mathbf{I}_{ij} - n_{ia}T^{-1}_{a\nu}n_{j\nu}, \tag{39}$$

where

$$T_{a\nu} = n_{ia}n_{i\nu}. \tag{40}$$

The dynamical projection operator is used to project forces to $3N - Q$ dimensional hypersurface. Therefore, they are locally perpendicular to the constraints.

The mid-step algorithm proposed by Hinch is for the case of mobility tensor (33) composed by four following substeps:

1. Generate unprojected random forces $\eta_i$ and unprojected forces $f_i$ at initial position $x^0_i$;
2. Construct projected random force $\eta^P_i = P_{ij}\eta_j$ and $f^P_i = P_{ij}f_j$;
3. Calculate midstep position $x^{1/2}_i = x^0_i + \dot{x}^0_i\Delta t/2$, where the mobility in the original configuration $\dot{x}^0_i$ is calculated via (37) and $\Delta t$ is the time step;
4. Calculate updated bead positions $x^1_i = x^0_i + \dot{x}^{1/2}_i\Delta t$, where $\dot{x}^{1/2}_i$ is evaluated with the deterministic and normal vectors from mid position, but with the same projected random force from initial configuration.

Mid-step algorithm uses the projection operator (38) that alongside with the mid position calculation minimizes the perturbations of constraints. Nevertheless, perturbations cannot be eliminated. Therefore, the MT has to be resized to fulfill the constraints. In such operation, the angles between $\vec{t_i}$ and $\vec{t_{i+1}}$, where $i = 1, ..., N - 1$ are conserved, the first bead of MT remains constant and MT regrows from the MTOC. Consequently, the bending energy of the MT remains unchanged.

### 3 Additional results

#### 3.1 Influence of random forces

Random forces acting on the MTs have small effects since the motion of the MTs is constrained. Figure 5a suggests that random forces perpendicular to the cell membrane(red) have no effect since microtubule curvature is given by the interplay of bending forces and the force of the membrane. The green forces, acting parallel to the segments of the MTs have also no impact, because the MT is attached to the MTOC, making it a part of a very massive structure. Consequently, just random forces depicted in purple in 5b can result in movement, making the random noise effectively one-dimensional. However, the MT is still rigid structure, random forces act in contradiction and filament is bound to MTOC, which seriously limits the movement of upper beads. Therefore, the influence of random noise can be expected to be negligible. Moreover, the capture-shrinkage mechanism fixates the MT on both sides, further minimizing the effect of the random force. In Figures 6a and 6b we can see that the repositioning curves are almost identical for the case of the capture-shrinkage mechanism. The figures 6c and 6d demonstrate that the developments of the number of attached dyneins also do not differ. During cortical sliding repositionings the effect of the random forces is also negligible 7.

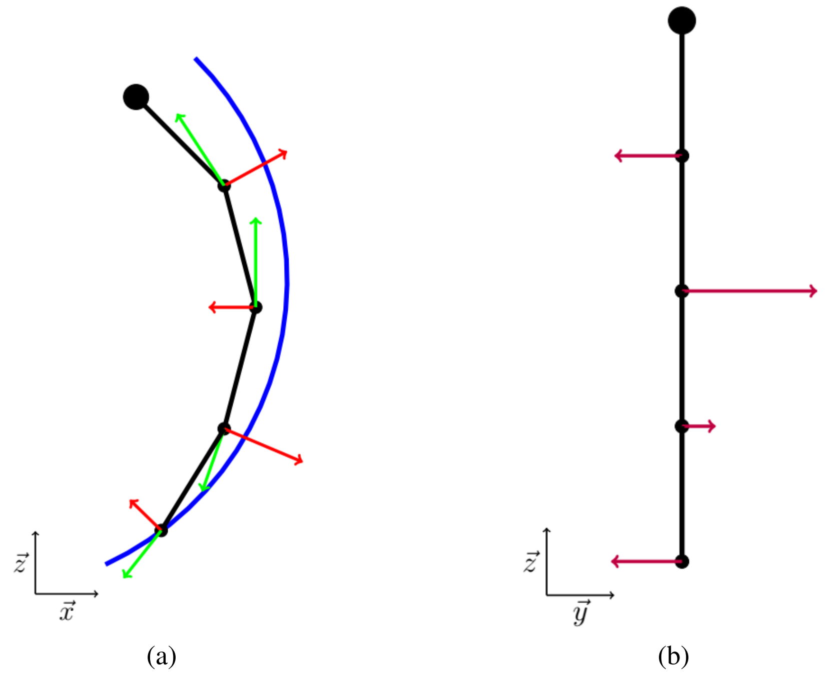


Figure 5: Sketch of random forces acting on microtubule from two perspectives. The big circle represents MTOC and smaller circles represent beads of microtubule. Blue arc depicts membrane of the cell. Red, green and purple lines representing random forces acting on every bead are perpendicular to each.

#### 3.2 Comparison of two cases with different number of microtubules

The cytoskeleton of $M_{\text{micro}} = 100$ MTs (examined in previous section) is compared with the cytoskeleton of $M_{\text{micro}} = 40$. We define $n_{\text{dm}}(t) = \frac{\tilde{N}_{\text{dm}}(t)}{M_{\text{micro}}}$ to examine the ratio of the attached dyneins and the number of the MTs in a cytoskeleton. Figure 8a depicts repolarization curves of two cytoskeletons for the case of the capture-shrinkage mechanism. The polarization exhibits a triphasic behavior for both cases.

We define $n_{\text{max}} = \max(n_{\text{dm}}(t))$. In Figure 8c it can be seen that $n_{\text{max}}$ is always bigger for the case of smaller cytoskeleton(caused by the small area of the center of IS and limited number of dynein). The Figures 8c and 8d explain the differences of speed in terms of the number of motors. When the area density is small, the smaller cytoskeleton is pulled with relatively higher force. As the concentration increases, the maximum speed is achieved($\rho_{\text{IS}} \sim 600\mu\text{m}^{-2}$). Subsequent increase of pulling force has no effect.

Figures 9a and 9b show the same trends for both cytoskeletons in the case of cortical sliding mechanism. We observe the three characteristics based on area density for both cytoskeleton. In both cases, $n_{\text{max}}$ rises at the beginning, it reaches its maximum when $\tilde{\rho}_{\text{IS}} \sim 200\mu\text{m}^{-2}$ and then it decreases swiftly and then steadily 9c. For smaller densities, the number of dyneins per MT are smaller for the bigger cytoskeleton. The situation is opposite when considering high area densities. Since the attached MTs aim in different directions, dyneins compete in the area of higher densities. As the number of the MT decreases, the pulling forces acting on individual filaments increase, leading to faster detachment. The MTOC speed increases when $\tilde{\rho}_{\text{IS}} < 200\mu\text{m}^{-2}$ and then it decreases. We can see in 9d that the speed decreases for both cases even when $n_{\text{max}}$ stays approximately the same, which is the consequence of dynein acting predominantly at the periphery.

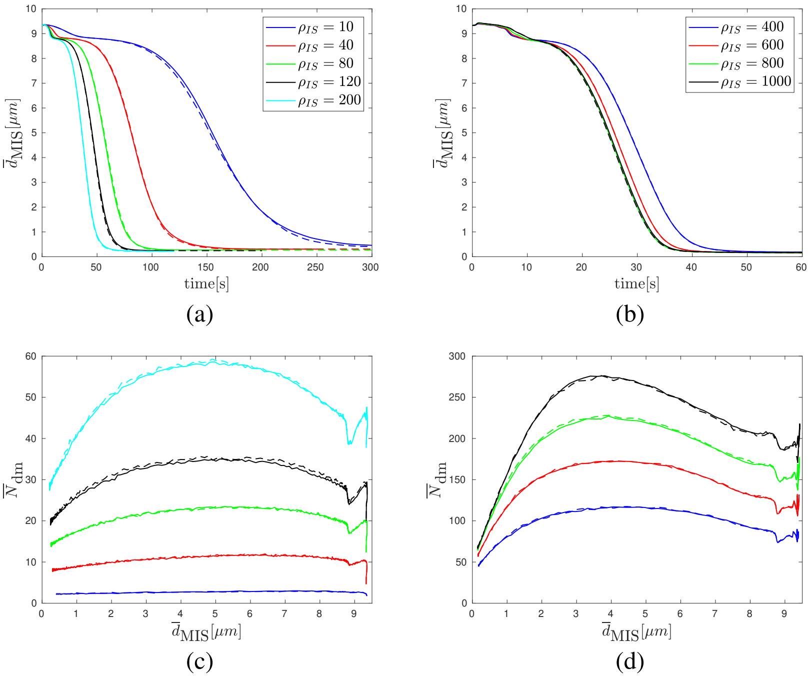


Figure 6: Repositioning curves under the influence of capture-shrinkage and random forces. Legends in (a), (b) apply for (c), (d), respectively. Dashed curves comprise random forces. (a)(b) Dependence of MTOC-IS distance $\bar{d}_{\text{MIS}}$ on time, (c)(d) Dependence of the number of dynein $\bar{N}_{\text{dm}}$ motors on MTOC-IS distance $\bar{d}_{\text{MIS}}$.

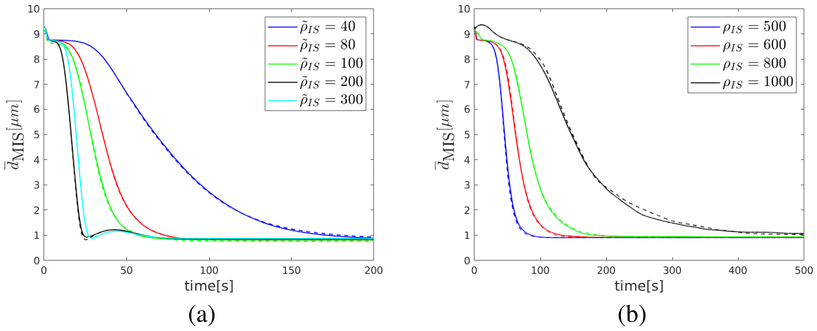


Figure 7: Repositioning curves under influence of cortical sliding and random forces. (a)(b) Dependence of average MTOC-IS distance $\bar{d}_{\text{MIS}}$ on time,

##### 3.3 Capture-shrinkage and cortical sliding combined

As can be seen in Figure 10a, addition of the small area density of capture-shrinkage dyneins in the center of IS causes substantial decrease of differences between times of polarization. Moreover, the three regimes of behavior based on the area density of cortical sliding dyneins is not observed in the presence of capture-shrinkage. Surely, the third regime presents a disadvantage, since the pulling force of dynein is wasted in unproductive competitions. Therefore, the synergy of two mechanisms proves once more to be highly effective, since it does not only removes the third regime, but also greatly reduces the times of repositioning when the area density of cortical sliding $\tilde{\rho}_{\text{IS}} < 100\mu\text{m}^{-2}$. Figure 10b depicts times of repositioning for different sets of combined mechanisms since the capture-shrinkage area density varies and cortical sliding density remains constant. We can see that the times of repositioning are in general shorter for the case of combined mechanisms. Moreover, even when the area densities correspond to second regime, the times of repositioning are comparable. Combined cases, however, have just $15\%$ of the number of dyneins. Additionally, we can notice that the increase of area densities when $\tilde{\rho}_{\text{IS}} > 500\mu\text{m}^{-2}$ presents no advantage since it causes slowing down of repositioning in the absence of capture-shrinkage and has no effect when the mechanisms are combined. Figure 10c shows that the attached dyneins are predominantly located on the periphery of IS even in the case when $\tilde{\rho}_{\text{IS}} > \rho_{\text{IS}}$.

#### 4 Commentary on modeling approaches

##### 4.1 Cytosim

Cytosim is widely accepted as an efficient tool for the simulations of cytoskeletal fibers (55). Although there are many similarities between the models, we decided not to use Cytosim. The first reason is our goal to examine the role of

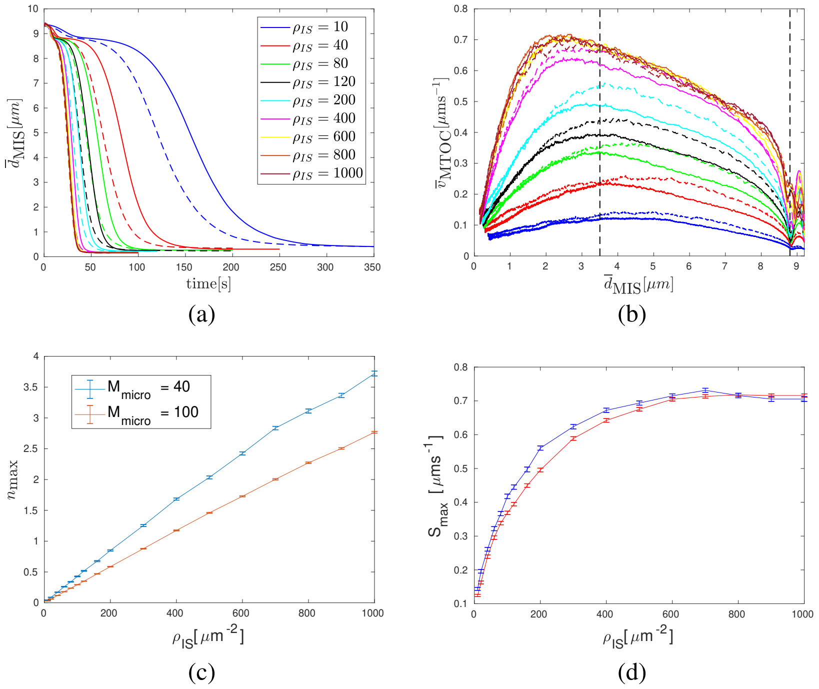


(a) (b)

(c) (d)

Figure 8: Capture-shrinkage mechanism for two cytoskeletons with different numbers of microtubules. Comparison of repolarization curves for cytoskeleton of $M_{\mathrm{micro}} = 100$(solid lines) and $M_{\mathrm{micro}} = 40$(dashed lines). (a) Dependence of average MTOC-IS distance $\bar{d}_{\mathrm{MIS}}$ on time. (b) Dependence of average MTOC speed $\bar{v}_{\mathrm{MTOC}}$ on $\bar{d}_{\mathrm{MIS}}$. (c) Dependence of the number of attached dyneins per microtubule $n_{\max}$ on area density $\rho_{\mathrm{IS}}$. (d) Dependence of the maximum speed $S_{\max}$ on $\rho_{\mathrm{IS}}$.

Brownian motion. The implicit integration used by Cytosim has a numerical error that could influence the precision of calculation in the absence of thermal noise. The second reason is the simplified calculation of the bending forces used by Cytosim. The advantage of such an approach, which enables to express the bending forces as a result of matrix-vector multiplication, is a great efficiency of calculation. Nevertheless, the procedure is valid only if the angles between subsequent segments remain small. This presents a drawback, since the angles between the segment increase as the radius of the cell decrease. More importantly, substantial curvature of the MTs can be expected during repositioning (56)(See 4.4). Moreover, the "reshaping" of the objects due to the numerical impressions is done to keep the center of mass constant. Since the rigidity of MTOC is an important part of our model, reshaping is done to keep the first the bead of the MT(therefore MTOC-MT forces) constant.

#### 4.2 Model using deterministic force

Kim and Maly (57) modeled the cortical sliding mechanism using the deterministic force. Although this model has various merits, we discourage of the modeling of the cortical sliding using a deterministic force since it leads to the contradiction with the experimental observable. Using the deterministic force leads to MTs stalk going through the center (57). This presents a drawback since the biological functions depend on the distribution of MTs. Subsequently, it is hard to see the biological motivation for the limiting of the MTOC movement on the surface of the nucleus, since the MTOC goes through radial movement (58). Moreover, it is questionable, if the deterministic force would be able of any action in the cell with less favorable shape. MTs, in general, do not copy the surface of the cell, since they form an arc touching the membrane of the cell just at the end(which is a consequence of energy minimalization). The flattening of cell and the fixation of MTOC on the nucleus permits close contact with the cell membrane, which is not achievable for a general shape.

#### 4.3 Movies

Movie 1 depicts MTOC repositioning just under the actions of capture-shrinkage mechanism $\rho_{textrmIS} = 100\mu\mathrm{m}^{-2}$. In the first seconds of repositioning, MTs passing through the center of IS attach to dynein. As the MTOC moves to the IS, MTs form a narrow stalk and the MT spindle opens. The transition between two phases takes place at the time $t \sin 55$s.

Movie 2 displays capture-shrinkage repositioning in the cell with smaller nucleus $r_{\mathrm{N}} = 3.3\mu\mathrm{m}$. the reduction of the nucleus enables to clearly see the change between two phases including the reduction of speed and the direction of MTOC motion.

Movie 3 shows the repositioning under the sole actions of cortical sliding with area density $\tilde{\rho}_{textrmIS} = 60\mu\mathrm{m}^{-2}$ belonging to the first regime. Majority of MTs attaches due to the relatively big area of IS. Subsequently, as MTOC approaches IS, MTs detach and the remaining filaments form a stalk. The speed of MTOC decreases at the end of

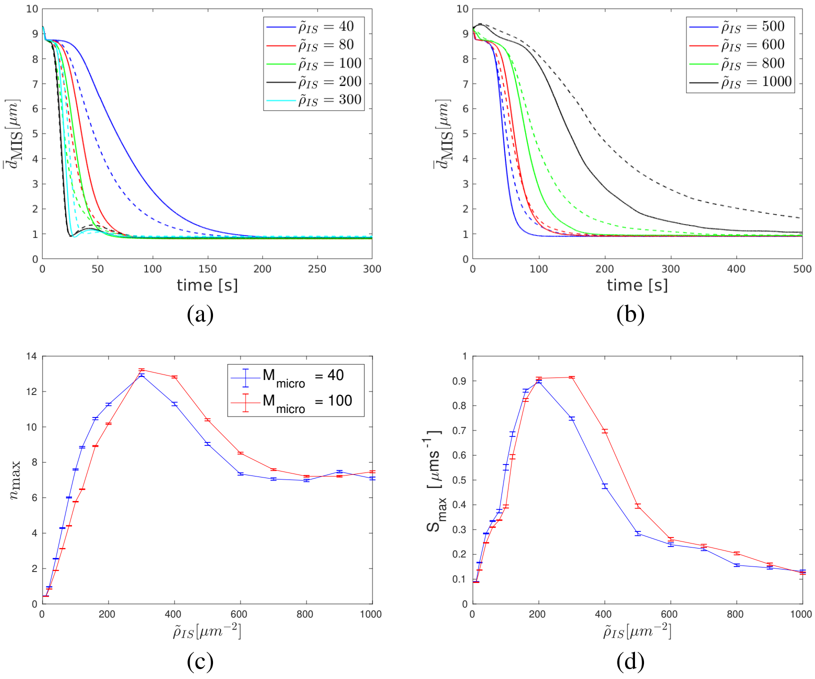


Figure 9: Cortical sliding mechanism for two cytoskeletons with different numbers of microtubules. Comparison of repolarization curves for cytoskeleton of $M_{\text{micro}} = 100$(solid lines) and $M_{\text{micro}} = 40$(dashed lines). (a)(b) Dependence of average MTOC-IS distance $\bar{d}_{\text{MIS}}$ on time. (c) Dependence of the number of attached dyneins per microtubule $n_{\max}$ on area density $\tilde{\rho}_{\text{IS}}$. (d) Dependence of the maximum speed $S_{\max}$ on $\tilde{\rho}_{\text{IS}}$.

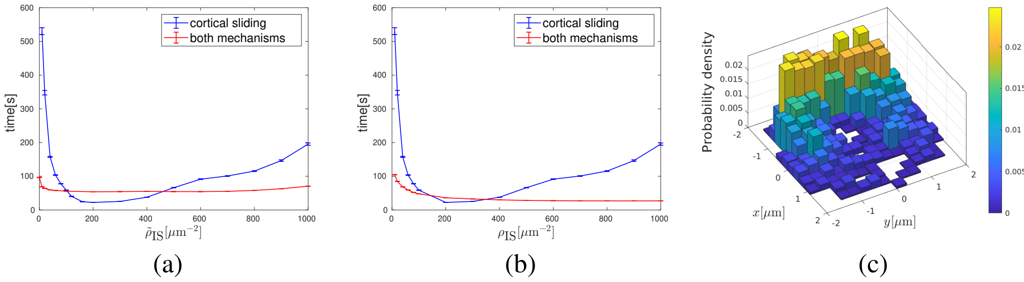


Figure 10: Combined mechanisms. (a)(b) Times of repositioning for the sole cortical sliding and combinations of two mechanisms. (a)In the combined case the cortical sliding area density $\tilde{\rho}_{\text{IS}}$ varies and capture-shrinkage area density is constant $\tilde{\rho}_{\text{IS}} = 60\mu\text{m}^{-2}$. (b) The capture-shrinkage area varies, cortical sliding area density is constant $\tilde{\rho}_{\text{IS}} = 20\mu\text{m}^{-2}$. (c) Two dimensional probability density of attached dynein $\tilde{\rho}_{\text{IS}} = 60\mu\text{m}^{-2}$, $\rho_{\text{IS}} = 20\mu\text{m}^{-2}$, $\bar{d}_{MIS} = 4.5\mu\text{m}$.

repositioning and MTOC does not recedes from the nucleus. It can be seen that the majority of motors is caught at the periphery of IS.

Movie 4 video depicts the second regime of cortical sliding $\tilde{\rho}_{\text{IS}} = 200\mu\text{m}^{-2}$. In comparison with the previous video, there are three major differences. Most notable is the speed of repositioning that is the consequence of formation of a very large stalk. Moreover, the MTOC passes the entire IS and,subsequently, returns to the center of IS.

Movie 5 video visualizes the repositioning in the third regime $\tilde{\rho}_{\text{IS}} = 1000\mu\text{m}^{-2}$. Due to the high number of dynein motors, it takes considerable time before the prevalent direction of MTOC motion is established. Subsequently, MTOC moves to the IS. The localization of the attached dynein motors is the most notable in the third regime.

Movies 6 and 7 are short and depicts the crucial difference between the sole capture-shrinkage and cortical sliding. In the sixth video, just the long MTs are attached in the center of IS. On the contrary, all MTs intersecting IS are caught in the combined case. Movies 7 displays that the MTs under the actions of cortical sliding are pulled to the center of IS.

Movie 8 shows the repositioning under the combined actions of both mechanisms, $\rho_{\text{IS}} = \tilde{\rho}_{\text{IS}} = 60\mu\text{m}^{-2}$. At the beginning, due to the actions of cortical sliding, large majority of MTs are caught in the center of IS(compare with Movie 7). Since the MTs originally sprout in different direction, just a fraction is able to form a stalk. The not-belonging to the stalk detach during the repositioning. Subsequently, they are caught by the cortical sliding, forming the arc patterns in the IS.

### References


1. Broedersz, C., and F. MacKintosh, 2014. Modeling semiflexible polymer networks. *Rev. Mod. Phys.* 86:995–1036.

2. Howard, J., 2001. Mechanics of Motor Proteins and the Cytoskeleton. Sinauer Associates is an imprint of Oxford University Press, Sunderland, Mass, new edition edition.

3. Leith, D., 1987. Drag on Nonspherical Objects. *Aerosol Science and Technology* 6:153–161.

4. Maccari, I., R. Zhao, M. Peglow, K. Schwarz, I. Hornak, M. Pasche, A. Quintana, M. Hoth, B. Qu, and H. Rieger, 2016. Cytoskeleton rotation relocates mitochondria to the immunological synapse and increases calcium signals. *Cell Calcium* 60:309–321.

5. Bereiter-Hahn, J., and M. Vöth, 1994. Dynamics of mitochondria in living cells: shape changes, dislocations, fusion, and fission of mitochondria. *Microsc. Res. Tech.* 27:198–219.

6. Jakobs, S., 2006. High resolution imaging of live mitochondria. *Biochimica et Biophysica Acta (BBA) - Molecular Cell Research* 1763:561–575.

7. Chaudhuri, A., 2016. Cell Biology by the Numbers. *Yale J Biol Med* 89:425–426.

8. 1966. An Atlas of Fine Structure. The Cell. Its Organelles and Inclusions. *Ann Intern Med* 64:968.

9. Jakobs, S., and C. A. Wurm, 2014. Super-resolution microscopy of mitochondria. *Current Opinion in Chemical Biology* 20:9–15.

10. Xu, H., W. Su, M. Cai, J. Jiang, X. Zeng, and H. Wang, 2013. The Asymmetrical Structure of Golgi Apparatus Membranes Revealed by In situ Atomic Force Microscope. *PLoS One* 8.

11. Ladinsky, M. S., D. N. Mastronarde, J. R. McIntosh, K. E. Howell, and L. A. Staehelin, 1999. Golgi Structure in Three Dimensions: Functional Insights from the Normal Rat Kidney Cell. *J Cell Biol* 144:1135–1149.

12. Day, K. J., L. A. Staehelin, and B. S. Glick, 2013. A Three-Stage Model of Golgi Structure and Function. *Histochem Cell Biol* 140:239–249.

13. Huang, S., and Y. Wang, 2017. Golgi structure formation, function, and post-translational modifications in mammalian cells. *F1000Res* 6.

14. Westrate, L. M., J. E. Lee, W. A. Prinz, and G. K. Voeltz, 2015. Form follows function: the importance of endoplasmic reticulum shape. *Annu. Rev. Biochem.* 84:791–811.

15. English, A. R., and G. K. Voeltz, 2013. Endoplasmic reticulum structure and interconnections with other organelles. *Cold Spring Harb Perspect Biol* 5:a013227.

16. English, A. R., N. Zurek, and G. K. Voeltz, 2009. Peripheral ER structure and function. *Curr. Opin. Cell Biol.* 21:596–602.

17. Shibata, Y., G. K. Voeltz, and T. A. Rapoport, 2006. Rough sheets and smooth tubules. *Cell* 126:435–439.

18. Hu, J., W. A. Prinz, and T. A. Rapoport, 2011. Weaving the Web of ER Tubules. *Cell* 147:1226–1231.

19. Gurel, P. S., A. L. Hatch, and H. N. Higgs, 2014. Connecting the Cytoskeleton to the Endoplasmic Reticulum and Golgi. *Curr. Biol.* 24:R660–R672.

20. Alberts, B., A. Johnson, J. Lewis, M. Raff, K. Roberts, and P. Walter, 2007. Molecular Biology of the Cell, 5th Edition. Garland Science, New York, 5th edition edition.

21. Goodenough, U. W., B. Gebhart, V. Mermall, D. R. Mitchell, and J. E. Heuser, 1987. High-pressure liquid chromatography fractionation of Chlamydomonas dynein extracts and characterization of inner-arm dynein subunits. *Journal of Molecular Biology* 194:481–494.

22. Gee, M. A., J. E. Heuser, and R. B. Vallee, 1997. An extended microtubule-binding structure within the dynein motor domain. *Nature* 390:636–639.

23. Goodenough, U., and J. Heuser, 1984. Structural comparison of purified dynein proteins with in situ dynein arms. *Journal of Molecular Biology* 180:1083–1118.

24. Schmidt, H., E. S. Gleave, and A. P. Carter, 2012. Insights into dynein motor domain function from a 3.3 Å crystal structure. *Nat Struct Mol Biol* 19:492–S1.

25. Leduc, C., O. Campàs, K. B. Zeldovich, A. Roux, P. Jolimaitre, L. Bourel-Bonnet, B. Goud, J.-F. Joanny, P. Bassereau, and J. Prost, 2004. Cooperative extraction of membrane nanotubes by molecular motors. *Proc. Natl. Acad. Sci. U.S.A.* 101:17096–17101.

26. Kamiya, N., T. Mashimo, Y. Takano, T. Kon, G. Kurisu, and H. Nakamura, 2016. Elastic properties of dynein motor domain obtained from all-atom molecular dynamics simulations. *Protein Eng Des Sel* 29:317–325.

27. Burgess, S. A., M. L. Walker, H. Sakakibara, P. J. Knight, and K. Oiwa, 2003. Dynein structure and power stroke. *Nature* 421:715–718.

28. Lindemann, C. B., and A. J. Hunt, 2003. Does axonemal dynein push, pull, or oscillate? *Cell Motil. Cytoskeleton* 56:237–244.

29. Sakakibara, H., H. Kojima, Y. Sakai, E. Katayama, and K. Oiwa, 1999. Inner-arm dynein c of Chlamydomonas flagella is a single-headed processive motor. *Nature* 400:586–590.

30. Sakakibara, H., and K. Oiwa, 2011. Molecular organization and force-generating mechanism of dynein. *The FEBS Journal* 278:2964–2979.

31. Gennerich, A., A. P. Carter, S. L. Reck-Peterson, and R. D. Vale, 2007. Force-Induced Bidirectional Stepping of Cytoplasmic Dynein. *Cell* 131:952–965.

32. Toba, S., T. M. Watanabe, L. Yamaguchi-Okimoto, Y. Y. Toyoshima, and H. Higuchi, 2006. Overlapping hand-over-hand mechanism of single molecular motility of cytoplasmic dynein. *PNAS* 103:5741–5745.

33. Mallik, R., D. Petrov, S. A. Lex, S. J. King, and S. P. Gross, 2005. Building complexity: an in vitro study of cytoplasmic dynein with in vivo implications. *Curr. Biol.* 15:2075–2085.

34. Mallik, R., B. C. Carter, S. A. Lex, S. J. King, and S. P. Gross, 2004. Cytoplasmic dynein functions as a gear in response to load. *Nature* 427:649–652.

35. Reck-Peterson, S. L., A. Yildiz, A. P. Carter, A. Gennerich, N. Zhang, and R. D. Vale, 2006. Single-Molecule Analysis of Dynein Processivity and Stepping Behavior. *Cell* 126:335–348.

36. Kural, C., H. Kim, S. Syed, G. Goshima, V. I. Gelfand, and P. R. Selvin, 2005. Kinesin and dynein move a peroxisome in vivo: a tug-of-war or coordinated movement? *Science* 308:1469–1472.

37. Torisawa, T., M. Ichikawa, A. Furuta, K. Saito, K. Oiwa, H. Kojima, Y. Y. Toyoshima, and K. Furuta, 2014. Autoinhibition and cooperative activation mechanisms of cytoplasmic dynein. *Nature Cell Biology* 16:1118–1124.

38. Müller, M. J. I., S. Klumpp, and R. Lipowsky, 2008. Tug-of-war as a cooperative mechanism for bidirectional cargo transport by molecular motors. *Proc. Natl. Acad. Sci. U.S.A.* 105:4609–4614.

39. King, S. J., and T. A. Schroer, 2000. Dynactin increases the processivity of the cytoplasmic dynein motor. *Nat Cell Biol* 2:20–24.

40. Nishiura, M., T. Kon, K. Shiroguchi, R. Ohkura, T. Shima, Y. Y. Toyoshima, and K. Sutoh, 2004. A single-headed recombinant fragment of Dictyostelium cytoplasmic dynein can drive the robust sliding of microtubules. *J. Biol. Chem.* 279:22799–22802.

41. Kon, T., M. Nishiura, R. Ohkura, Y. Y. Toyoshima, and K. Sutoh, 2004. Distinct Functions of Nucleotide-Binding/Hydrolysis Sites in the Four AAA Modules of Cytoplasmic Dynein. *Biochemistry* 43:11266–11274.

42. Cho, C., S. L. Reck-Peterson, and R. D. Vale, 2008. Regulatory ATPase sites of cytoplasmic dynein affect processivity and force generation. *J. Biol. Chem.* 283:25839–25845.

43. Kikushima, K., T. Yagi, and R. Kamiya, 2004. Slow ADP-dependent acceleration of microtubule translocation produced by an axonemal dynein. *FEBS Letters* 563:119–122.

44. Walter, W. J., B. Brenner, and W. Steffen, 2010. Cytoplasmic dynein is not a conventional processive motor. *J. Struct. Biol.* 170:266–269.

45. Belyy, V., M. A. Schlager, H. Foster, A. E. Reimer, A. P. Carter, and A. Yildiz, 2016. The mammalian dynein-dynactin complex is a strong opponent to kinesin in a tug-of-war competition. *Nat. Cell Biol.* 18:1018–1024.

46. Ikuta, J., N. K. Kamisetty, H. Shintaku, H. Kotera, T. Kon, and R. Yokokawa, 2014. Tug-of-war of microtubule filaments at the boundary of a kinesin- and dynein-patterned surface. *Scientific Reports* 4:5281.

47. Kunwar, A., S. K. Tripathy, J. Xu, M. K. Mattson, P. Anand, R. Sigua, M. Vershinin, R. J. McKenney, C. C. Yu, A. Mogilner, and S. P. Gross, 2011. Mechanical stochastic tug-of-war models cannot explain bidirectional lipid-droplet transport. *Proc. Natl. Acad. Sci. U.S.A.* 108:18960–18965.

48. Klein, S., C. Appert-Rolland, and L. Santen, 2015. Motility states in bidirectional cargo transport. *EPL* 111:68005.

49. Montesi, A., D. C. Morse, and M. Pasquali, 2005. Brownian dynamics algorithm for bead-rod semiflexible chain with anisotropic friction. *J. Chem. Phys.* 122:084903.

50. Fixman, M., 1978. Simulation of polymer dynamics. I. General theory. *J. Chem. Phys.* 69:1527–1537.

51. Hinch, E. J., 1994. Brownian motion with stiff bonds and rigid constraints. *Journal of Fluid Mechanics* 271:219–234.

52. Grassia, P. S., E. J. Hinch, and L. C. Nitsche, 1995. Computer simulations of Brownian motion of complex systems. *Journal of Fluid Mechanics* 282:373–403.

53. Grassia, P., and E. J. Hinch, 1996. Computer simulations of polymer chain relaxation via Brownian motion. *Journal of Fluid Mechanics* 308:255–288.

54. Pasquali, M., and D. C. Morse, 2002. An efficient algorithm for metric correction forces in simulations of linear polymers with constrained bond lengths. *The Journal of Chemical Physics* 116:1834–1838.

55. Nedelec, F., and D. Foethke, 2007. Collective Langevin dynamics of flexible cytoskeletal fibers. *New J. Phys.* 9:427–427.

56. Kuhn, J. R., and M. Poenie, 2002. Dynamic Polarization of the Microtubule Cytoskeleton during CTL-Mediated Killing. *Immunity* 16:111–121.

57. Kim, M. J., and I. V. Maly, 2009. Deterministic Mechanical Model of T-Killer Cell Polarization Reproduces the Wandering of Aim between Simultaneously Engaged Targets. *PLOS Computational Biology* 5:e1000260.

58. Yi, J., X. Wu, A. H. Chung, J. K. Chen, T. M. Kapoor, and J. A. Hammer, 2013. Centrosome repositioning in T cells is biphasic and driven by microtubule end-on capture-shrinkage. *J Cell Biol* 202:779–792.
